# A 37-million-particle dataset from over 250 experiments to accelerate data-driven cryo-EM analysis

**DOI:** 10.64898/2026.04.29.720997

**Authors:** Andreas Zamanos, Fotis L. Kyrilis, Panagiotis Koromilas, Panagiotis L. Kastritis, Yannis Panagakis

**Affiliations:** Department of Informatics and Telecommunications, National and Kapodistrian University of Athens, 16122 Athens, Greece; Archimedes Unit, Athena Research Center, 15125 Marousi, Greece; Institute of Chemical Biology, National Hellenic Research Foundation, 11635 Athens, Greece; Department of Integrative Structural Biochemistry, Institute of Biochemistry and Biotechnology, Martin Luther University Halle-Wittenberg, 06120 Halle/Saale, Germany; Biozentrum, Martin Luther University Halle-Wittenberg, 06120 Halle/Saale, Germany; Interdisciplinary Research Center HALOmem, Charles Tanford Protein Center, Martin Luther University Halle-Wittenberg, 06120 Halle/Saale, Germany.∗

## Abstract

Cryogenic Electron Microscopy (cryo-EM) has revolutionized structural biology by enabling near-atomic-resolution structure determination of biological macromolecules. Central to cryo-EM analysis are particles, namely 2D projections of biomolecules extracted from micrographs, which serve as the primary input for 3D reconstruction. While data-driven methods have transformed other scientific domains, their impact on cryo-EM remains limited because existing particle datasets are too small, too narrow in protein diversity, and lack rich per-particle annotations. We introduce cryoPANDA (cryo-EM Particles ANnotated DAtaset), comprising over 37 million annotated particles from 252 experiments spanning a wide range of protein types, more than 10-fold larger than prior collections. Each particle is accompanied by detailed annotations covering acquisition, classification, and reconstruction metadata, alongside the corresponding 3D electrostatic potential map, the published EMDB map, and, where available, the PDB model. We validate cryoPANDA in two ways: first, by reconstructing hundreds of distinct high-resolution cryo-EM maps; and second, by training a DINOv2 foundation model and evaluating its learned representations on micrograph segmentation, particle picking, and particle clustering.

## Background & Summary

Cryo-electron microscopy (cryo-EM) has transformed structural biology by allowing the visualization of biological macromolecules at near-atomic resolution, including many systems that are difficult to study using other structural methods such as NMR spectroscopy or X-ray crystallography [1–4]. In a typical single-particle cryo-EM experiment, a purified macromolecular sample is rapidly frozen in a thin layer of vitreous ice, preserving its native conformation while minimizing radiation damage. The frozen specimen is then imaged using an electron microscope, producing two-dimensional projections of individual molecules in images known as *micrographs*. Each micrograph contains many randomly oriented copies of the target molecule, referred to as *particles*. These particles are identified (picked) from the micrographs and used to reconstruct a three-dimensional (3D) Coulomb (electrostatic) potential map.

At sufficiently high resolution, an atomic model can be built and refined into the reconstructed map [5–7]. Such models can provide fundamental insight into molecular mechanisms [8] and support applications in drug discovery and biotechnology [9]. Despite the transformative impact of cryo-EM, recognized by the 2017 Nobel Prize in Chemistry [10], cryo-EM data analysis remains challenging because of the intrinsic complexities of both the imaging process and the biological samples. These challenges include the extremely low signal-to-noise ratio of particle images, imaging artifacts, sample heterogeneity, beam-induced motion, and conformational flexibility [11, 12].

### From Sample to Structure: An Overview of the Cryo-EM Workflow

A typical single-particle cryo-EM experiment proceeds through several stages, summarized in Figure 1. The process begins with **sample and grid preparation**: the protein of interest is purified and applied to an EM grid, where, in a method called vitrification, excess solution is blotted away and the grid is rapidly plunged into liquid ethane to embed the specimen in a thin layer of amorphous (vitreous) ice, preserving its native conformation [13].

**Figure 1:**
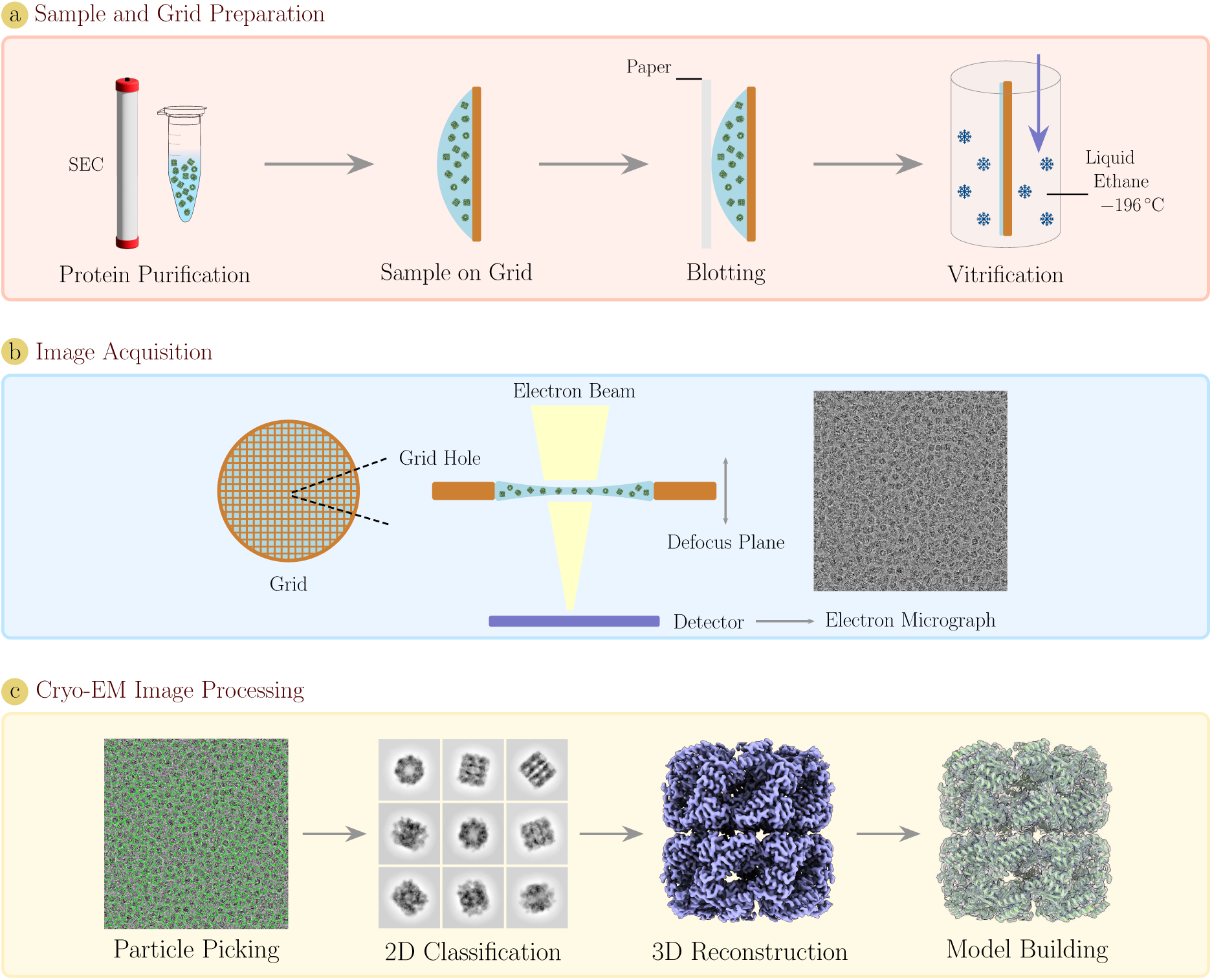
Overview of a standard cryo-EM workflow. Schematically, (a) illustrates the sample and grid preparation steps, (b) depicts the image acquisition process, and (c) outlines key image processing and analysis steps.

The frozen grid is then inserted into the electron microscope for **data acquisition**. For each selected grid hole, a dose-fractionated series of images is recorded as movie frames [9]. These frames are subsequently aligned and summed during **motion correction** to compensate for beam-induced specimen movement, yielding a single motion-corrected micrograph [14]. Because unstained biological specimens produce little contrast when imaged in focus, data are intentionally acquired under defocus, which introduces oscillations in the contrast transfer function (CTF) and masks high-resolution information. **CTF estimation** recovers these parameters from each micrograph, allowing their effects to be corrected in downstream processing [15].

With fully processed micrographs, the next critical step is **particle picking**: individual projections are detected and extracted from each micrograph. The quality of the resulting particle set strongly influences the attainable resolution of the final reconstruction [16]. Picked particles are then aligned and grouped during **2D classification**, which serves to discard false positives and low-quality images while organizing particles by viewing orientation — a process typically repeated iteratively under manual curation. The curated particle set is used for **3D reconstruction and refinement**, in which particle orientations are determined and iteratively updated to produce a three-dimensional electrostatic potential map of the protein. Finally, if the reconstruction reaches sufficient resolution, an atomic model can be built into the map, either by fitting and refining an existing model or by *de novo* placing of atoms.

Throughout this pipeline, particles are the central data unit: they are the direct output of picking, the input to classification and reconstruction, and ultimately determine the quality of the final structure. It is this pivotal role that motivates the construction of a large-scale, richly annotated particle dataset.

### The Data Bottleneck in Cryo-EM Methods Development

Machine learning has been widely adopted across cryo-EM analysis tasks over the past decade, including **particle picking** [17–32], **3D reconstruction** [33–41], **electrostatic potential map refinement** [42–44], and **atomic model building** [45–52]. Several of these methods have been successfully applied in real cryo-EM workflows by practitioners [53–55]. Despite this progress, most methods are either *trained independently for each experiment* or rely on *small, task-specific datasets*, limiting their ability to learn broadly transferable representations across the diversity of cryo-EM data. This has motivated a growing interest in **foundation models** — large neural networks trained on massive datasets to learn general-purpose representations that can be adapted to downstream tasks with minimal fine-tuning or in zero-shot settings.

Since 2024, foundation models have emerged for cryo-EM, trained on different data modalities: *micrographs* [56], *particles* [57, 58], and *electrostatic potential maps* [59, 60]. At the particle level — the modality most directly relevant to the cryo-EM processing pipeline — progress has been hindered by data availability: Cryo-IEF [57] relies on a *closed-source* dataset and demonstrates several key applications on simulated particles, while CryoFastAR [58] resorts to *simulated training data* due to the absence of large-scale, high-quality annotated experimental particle data, leading to performance degradation on real data. More broadly, a common prerequisite for a successful foundation model is the availability of a large and well-constructed dataset. The lack of a freely accessible, large-scale, and diverse experimental particle dataset therefore remains **a major bottleneck** for developing foundation models for cryo-EM analysis.

Two large curated cryo-EM datasets currently exist: CryoPPP [61] and CryoCRAB [62]. CryoPPP provides particle-picking annotations for micrographs from 34 cryo-EM experiments, comprising 9,893 micrographs and positional annotations for approximately 2.7 million particles; however, it is **limited in particle-level scale (L1)**, **spans few experiments with little diversity across protein types (L2)**, and **does not provide rich per-particle metadata** beyond picking coordinates **(L3)**. CryoCRAB aggregates data from 746 EMPIAR datasets, collecting 152,385 raw movies, but **provides annotations only at the micrograph level**. These resources are valuable, but neither offers a large, diverse corpus of particle images with rich per-particle annotations — the data most suited for down-stream cryo-EM analysis steps and for training particle-level foundation models. Without such a unified and comprehensive resource, current data-driven methods fail to generalize across the vast landscape of biological structures, requiring retraining and extra data for every new experimental setup **(L4)**. Beyond model development, cryoPANDA is designed to give cryo-EM practitioners direct access to particles, analysis metadata, 3D reconstructions, and atomic models for proteins or complexes of interest. A comparison between the three dataset is presented in Table 1.

**Table 1:** Comparison of CryoPPP, CryoCRAB, and cryoPANDA in terms of the number of experiments, micrographs, and particles. Bold indicates the highest value and underline the second highest. The asterisk on cryoPANDA micrographs indicates that these are not provided with the dataset but can be freely retrieved from EMPIAR.

|  | CryoPPP | CryoCRAB | cryoPANDA |
| --- | --- | --- | --- |
| Experiments | 34 | **756** | 252 |
| Micrographs | 9,893 | **152,385** | 103,457* |
| Particles | <u>2,700,000</u> | - | **37,623,123** |

Motivated by these limitations, we introduce cryoPANDA, an open, large-scale, and diverse particle dataset that **addresses all four gaps**. cryoPANDA contains more than **37 million** curated particle images from **252 experiments**, of which 247 are sourced from the Electron Microscopy Public Image Archive (EMPIAR) [63] and 5 from in-house experiments [64–66], spanning **16 functional protein classes** and totaling over **15 TB (L1, L2)**. Each particle is provided as an image with **detailed per-particle annotations** covering acquisition, classification statistics, and 3D reconstruction-related metadata **(L3)**. To facilitate efficient training and data access, we distribute particle images in HDF5 format, enabling high-throughput sampling and reducing I/O bottlenecks during model training. Finally, we demonstrate that cryoPANDA serves as the basis for overcoming **(L4)**; by pre-training a DINOv2 model on this vast data corpus, we show that the resulting representations excel at downstream tasks — including micrograph segmentation, particle picking, and particle clustering — without the need for any supervision or task-specific fine-tuning.

## Methods

### Data Acquisition

The design of cryoPANDA aims to achieve balanced coverage in terms of particle counts, experimental sources, and protein diversity. Our primary data source is EMPIAR [63]. We first select entries that directly provide picked particles; at the time of collection, EMPIAR contained 208 such entries. To complement these, we also include entries providing processed, motion-corrected micrographs, for which EMPIAR listed 287 entries. In total, we examined 495 unique EMPIAR entries.

### Filtering of Experiments

To ensure that our dataset is well balanced and does not repeatedly include similar protein types, we prune redundant entries. To this end, we retrieve 13,626 amino-acid sequences associated with the entries and perform pairwise sequence alignment between all distinct chains. Amino-acid sequence information is available for 476 of the 495 entries; the remaining 19 are excluded from this analysis. Based on a sequence similarity greater than 30%, we group the entries into 204 distinct clusters, which range in size from one to multiple entries. From each cluster, we select up to four representative EMPIAR entries to include in our dataset. This selection also involves manual curation to prioritize EMPIAR entries with well-documented experiments and high-quality data, while excluding entries with poor data quality or missing essential information. In total, 236 EMPIAR experiments are retained, of which 132 provide picked particles and 104 provide preprocessed single-frame micrographs. In addition, we include picked particles from 5 in-house cryo-EM experiments and 10 EMPIAR entries provided by CryoPPP, resulting in 252 distinct cryo-EM experiments. Note that one EMPIAR entry with picked particles contributed two distinct experiments.

### Metadata collection

Metadata are collected for each EMPIAR entry and its corresponding EMD entries from EMPIAR and EMDataResource [67], respectively. These metadata include pixel size, image dimensions (micrographs or particles), reconstruction resolution, symmetry, acceleration voltage, spherical aberration, electron dose, and other related parameters. Where a Protein Data Bank (PDB) [68] molecular model is associated with an EMPIAR entry, we compute the minimum and maximum diameters of the atomic model, in order to use this information as input parameters for particle picking.

### Data Collection

Once the EMPIAR entries are finalized and their metadata collected, we proceed with data acquisition. To maintain a balanced dataset, we set an initial target of approximately 300,000 particles per experiment, informed by prior cryo-EM studies that achieved sub-2 Å reconstructions from datasets of comparable size [69]. Because the final number of usable particles varies across experiments, we summarize the particle counts for all experiments in Figure S1. For entries with micrographs, we randomly sample 50 micrographs from each of the 104 EMPIAR experiments and perform particle picking to estimate the total number of micrographs required, imposing an upper limit of 1,000 micrographs per experiment. When author-provided particle coordinates are available for micrograph entries, we prioritize those over our own particle picking. For EMPIAR entries that provide picked particles directly, we estimate the number of particle files required to reach the target. In many cases, the target cannot be reached, which explains the discrepancy between the theoretical upper bound of approximately 75 million particles (252 *×* 300,000) and the final dataset size of more than 37 million particles. Once the final set of EMPIAR files is determined, we download all required data, amounting to roughly 29 TB distributed across more than 800,000 files.

### Data Analysis

Cryo-EM analysis is performed using cryoSPARC v4.6 [70]; an overview of the analysis is shown in Figure 2. We follow two general workflows, one for micrograph-based analysis and one for particle-based analysis, with additional steps applied when required for specific experiments.

**Figure 2:**
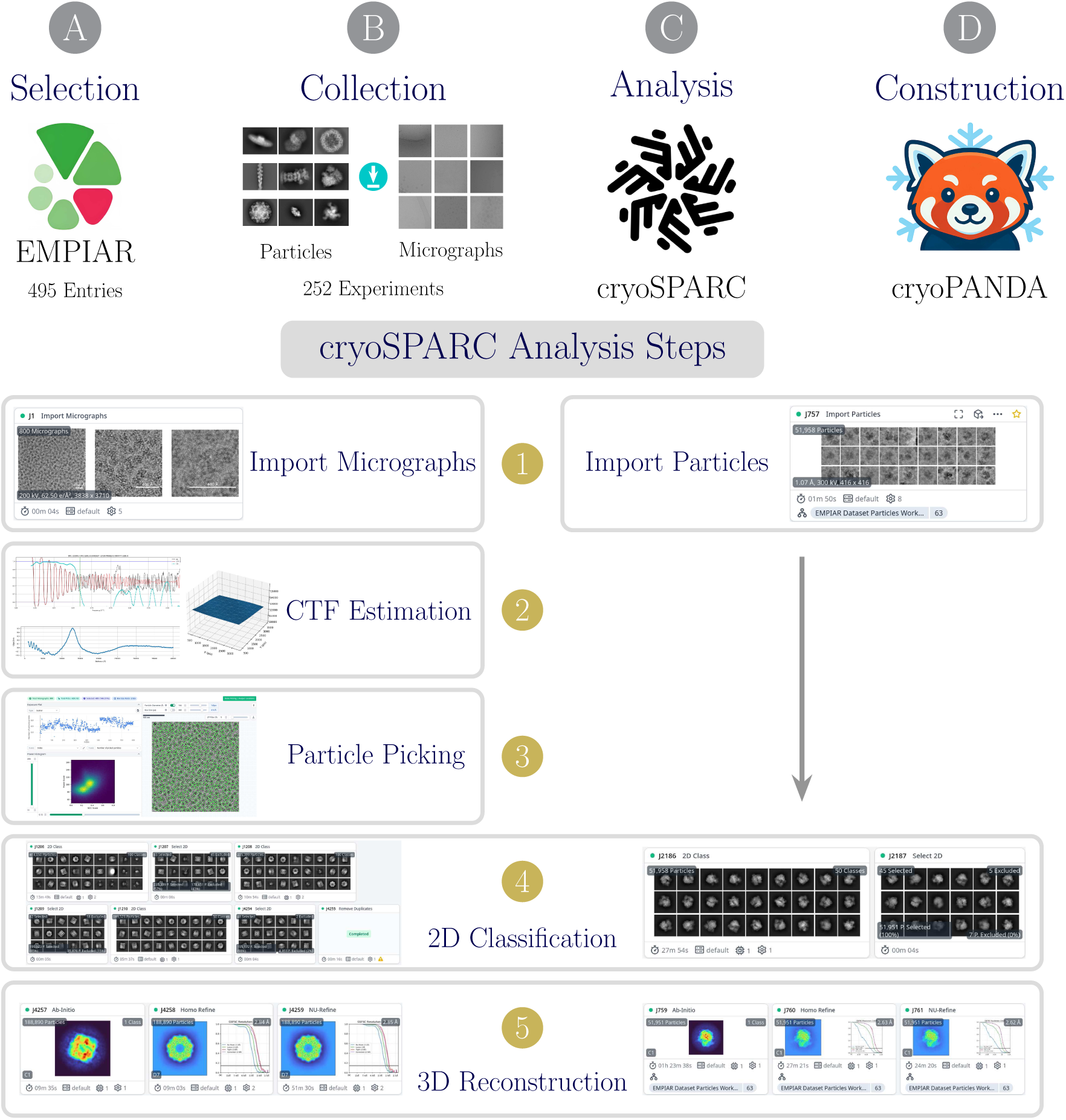
Overview of the cryoPANDA creation pipeline. (A) Selection of EMPIAR experiments. (B) Collection of particles and motion-corrected micrographs. (C) Analysis of the collected data. (D) Construction of the cryoPANDA dataset. The analysis steps are shown in detail for both particles and micrographs, from importing into cryoSPARC to 3D reconstruction.

### Micrograph Analysis

For micrograph analysis, we first **(1)** import the data and **(2)** perform Patch CTF Estimation (1) to determine the defocus landscape and astigmatism of each micrograph. Once CTF Estimation is completed, we proceed to **(3)** particle picking, using either the Blob Picker method or importing particle coordinates provided by the authors when available. After collecting the initial set of picked particles, we carry out three rounds of **(4)** 2D classification and selection. In each round, excluded particles are reclassified to recover any falsely discarded classes containing valid particles. At the end of this process, we remove potential duplicate particles using the minimum estimated particle diameter as a distance cut-off.

The Contrast Transfer Function (CTF) parameters estimated during the initial processing are defined by the phase-contrast approximation:

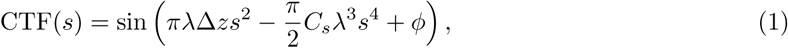

where *s* is the spatial frequency (in Å*^−^*^1^), *λ* is the electron wavelength, Δ*z* is the defocus (underfocus is positive), *C_s_* is the spherical aberration constant of the microscope, and *ϕ* is the total phase shift (e.g., due to amplitude contrast and, when present, a phase plate).

Once the particle set is finalized, we re-extract and restack the particles into MRC files containing 1,000 particles each. On these restacked particles, we perform a final 2D classification with 50 classes, from which we gather the clustering annotations for each particle. Finally, the restacked particles are used to **(5)** generate a 3D reconstruction of each experiment, typically following the sequence of *Ab Initio* Reconstruction, Homogeneous Refinement, and Non-Uniform Refinement, applying the appropriate symmetry during the latter two steps. The best map is selected based on the reported cryoSPARC resolution and visual agreement with the published electrostatic potential map.

### Particle Analysis

For the particle analysis, the workflow differs only in steps **(2)** and **(3)**, which are not required when picked particles are already available and can be imported directly into cryoSPARC. In addition, in step **(4)** we perform only a single round of 2D classification, used solely to exclude low-quality particles or residual noise left by the original picking.

### Analysis Results

In total, 1,710 and 3,100 individual cryoSPARC jobs are completed for particle-based and micrograph-based experiments, respectively. For experiments in which the *Ab Initio* Reconstruction fails to produce a good electrostatic potential map, we perform additional rounds of Homogeneous and Non-Uniform Refinement, as well as local CTF refinement. In cases where reconstruction cannot be obtained, we resort to initializing the 3D map using the published map. This process is necessary for 44 out of 252 experiments. Note that the maximum box size in our dataset is set to 448 pixels; this is primarily done to keep the total size of the cryoPANDA within practical limits. Box sizes exceeding this limit are downsampled to 448 pixels, either at the re-extraction step for experiments with micrographs or after **(1)** particle import for experiments with picked particles. Lastly, when the handedness of a reconstruction is incorrect, we flip the electrostatic potential map and rerun Homogeneous and Non-Uniform Refinement.

Out of the 252 experiments analyzed, we do not obtain a satisfactory 3D reconstruction for 38 experiments. For these cases we do not provide a 3D reconstruction generated by our pipeline. Nevertheless, particles from these experiments are retained in the final dataset, as they remain useful for particle-level analysis.

### Dataset Construction

Once the analysis is complete, we collect and organize the data. First, we record the job numbers corresponding to all steps described in Data Analysis for each experiment. We then transfer all required outputs from the cryoSPARC directories to our collection directory.

### Dataset Structure

The dataset is organized such that one directory is assigned to each experiment. Within each directory, the restacked particles (in batches of 1,000) are stored in a particles subdirectory, while the particles_information subdirectory contains metadata about the restack process, 2D classification, and 3D reconstruction of the particles. For each experiment, we also include the cryoPANDA reconstructed map, the published reconstructed map, the corresponding atomic model from the PDB when available, an annotation spreadsheet, and a particles.star file containing all particle metadata, which can be directly imported into analysis software such as cryoSPARC [70] or RELION [71].

### Data Annotations

Per-particle annotations are extensive and include: **(1)** general experimental information such as original and particle image pixel size (these differ only in the case of downsampling), particle box size, acceleration voltage, spherical aberration and electron dose; **(2)** protein-level information such as maximum molecular diameter in Ångstroms, symmetry, functional protein class, molecular weight, number of subunits and the presence of DNA or RNA in the structure; **(3)** CTF parameters, including defocus *U* and *V* (defoci along the two orthogonal principal astigmatism axes in the image plane), and the astigmatism angle; **(4)** micrograph-related information for experiments starting from micrographs, including micrograph pixel size, coordinates of picked particles, the normalized cross-correlation (NCC) score (which measures how well the particle aligns with the Gaussian during blob picking), and the micrograph shape and name; **(5)** 2D-classification metadata, including the particle’s class, the alignment resolution of that class, the class effective sample size (ESS) (2), and the effective classes assigned (ECA) score (3); **(6)** 3D-reconstruction metadata, including Euler angles (rotation, tilt, and psi), in-plane translations, the 3D alignment error with respect to the reconstructed map, and the final resolution of our electrostatic potential map; and **(7)** miscellaneous information such as associated EMD and PDB accession numbers and the resolution of the published map.

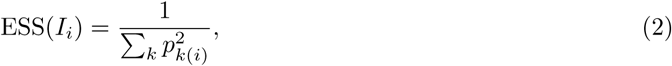

where *I_i_*is the *i*-th particle, *k* is the class index, and *p_k_*_(*i*)_ is the probability that *I_i_*belongs to class *k*.

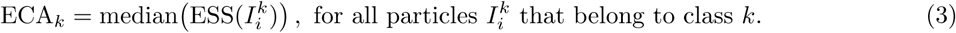

## Data Records

The cryoPANDA dataset is available at ScienceDB [72] under the DOI https://doi.org/10.57760/sciencedb.27164. The dataset contains 37,623,123 cryo-EM particles from 252 distinct experiments [53, 64–66, 73–285], most of which are from EMPIAR, and totals approximately 15.1 TB. A graphical representation of the dataset structure is presented in Figure 3; directories are shown in gray boxes and files in orange. The data are arranged in separate directories, one per cryo-EM experiment. Each experiment directory contains two subdirectories: particles, which stores restacked particles in batches of 1,000, and particles_information, which contains metadata related to restack, 2D classification, and 3D reconstruction jobs. These jobs are further organized into dedicated subdirectories and include output files from cryoSPARC, such as .png images and metadata files in .cs, .mrc, and .star formats. In addition to these two subdirectories, the main experiment directory contains (i) particles.star, which provides particle annotations required for importing particles into software packages such as cryoSPARC [70] or RELION [71], (ii) extended per-particle annotations in .xlsx format, (iii) the reconstructed electrostatic potential map produced by cryoPANDA in .mrc format, (iv) the corresponding EMDB map in .map format, and, where available, (v) an associated PDB atomic model in .cif format.

**Figure 3:**
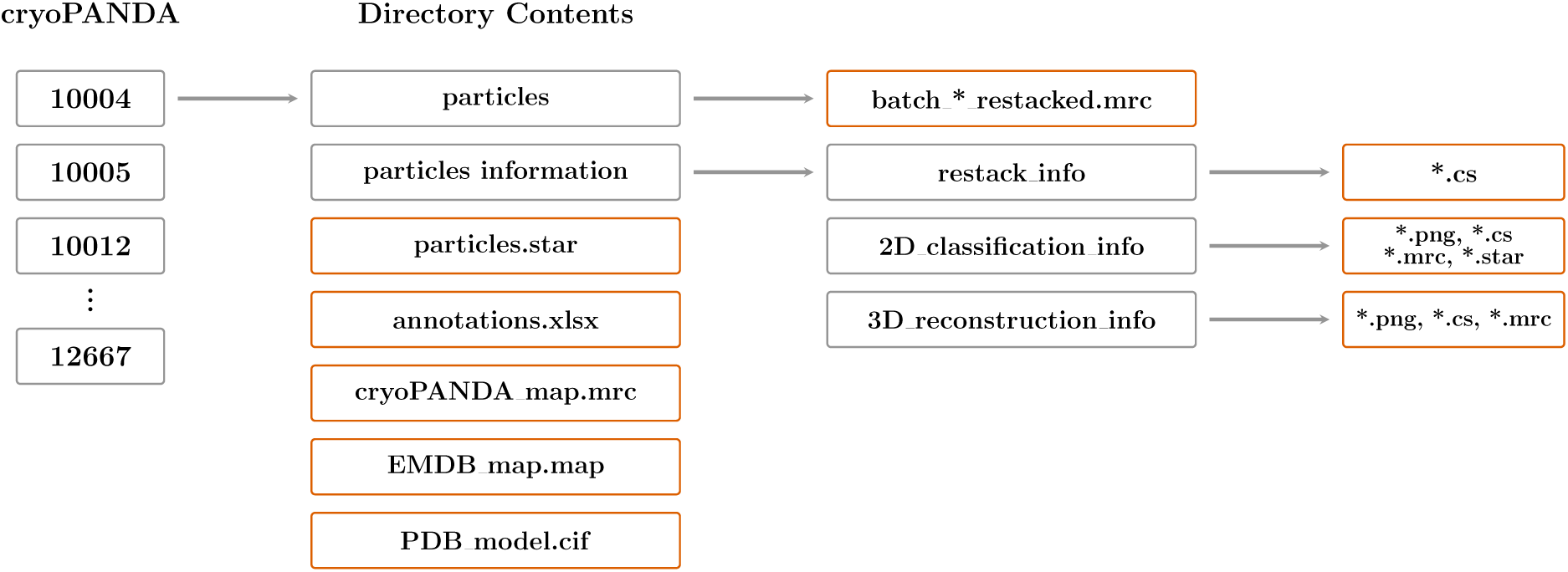
Directory structure of cryoPANDA. Black rectangular boxes denote directories, and orange boxes denote files. The hierarchy is organized by EMPIAR experiment, with each directory containing particles, associated metadata, 3D reconstructions, annotations, and, when available, the atomic model.

The cryoPANDA dataset is also provided in HDF5 format, chunked by experiment into 252 separate .h5 files. Within each HDF5 file, particle images and per-particle annotations can be accessed under the keys ’particles’ and ’annotations’, respectively. All particle images are resampled to 224 *×* 224 pixels, stored as uint8, and normalized per image to the [0–255] range. Combined with the smaller total size (1.8 TB) and the ease of distribution, this format significantly improves sampling efficiency during the training of machine-learning models.

### Data Overview

We provide a brief summary of the statistical distributions in the dataset. Figure 4 shows the molecular weight distribution of experiments and the protein classification distribution of particles in cryoPANDA. For clarity, we split the molecular weight into two (a and b) histograms at 800 kDa and limit the displayed range to 4000 kDa. The mean molecular weight is approximately 600 kDa (minimum 21 kDa, maximum 200,000 kDa); representative structures are shown above the histograms. In Figure 4c, we show the distribution of particles per protein class. These classes are defined based on cellular function rather than tertiary structure. Of the 16 classes, 10 have over 0.5 normalized particle count, suggesting a well-balanced and representative dataset across functional categories. Finally, Figure 5 summarizes the per-particle distributions of the following annotations: (a) EMDB reconstruction resolution, (b) pixel size, (c) symmetry, (d) defocus value, (e) maximum particle diameter, and (f) particle image size. Across the dataset, the mean reconstruction resolution is 3 Å, the mean pixel size is approximately 1 Å, and C1 is the most frequent symmetry class followed by C2. Defocus values range from 0 to 4 *µ*m (mean 1.7 *µ*m), maximum particle diameters are centered around 120 Å, and boxed particle sizes range from 128 to 448 pixels (mean 320 pixels).

**Figure 4:**
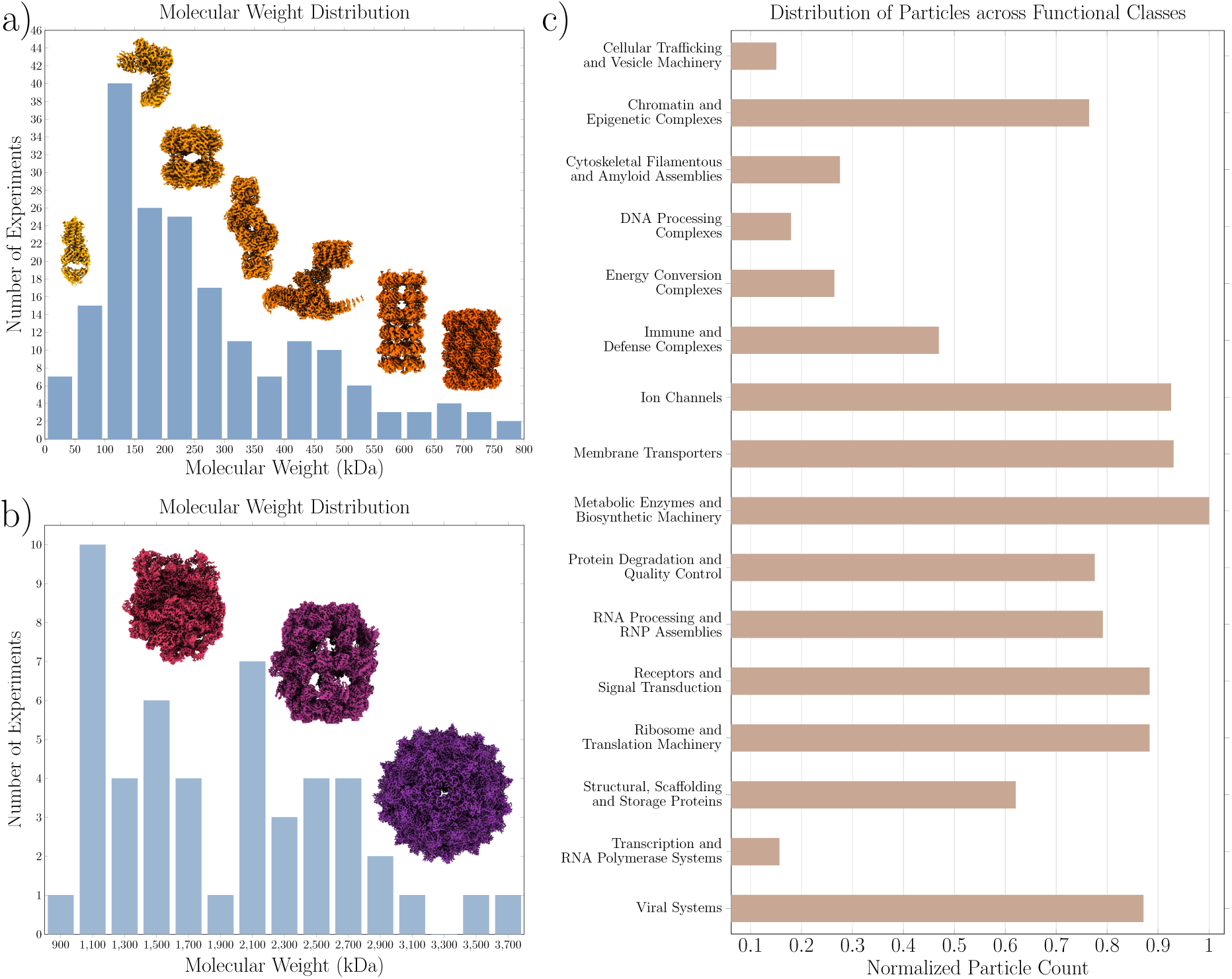
Molecular weight and protein classification distributions. Subplots (a) and (b) show the molecular weight distribution per experiment, split into two histograms at 800 kDa and limited to 4000 kDa. Subplot (c) shows the protein classification distribution by normalized particle count across 16 function-based classes.

**Figure 5:**
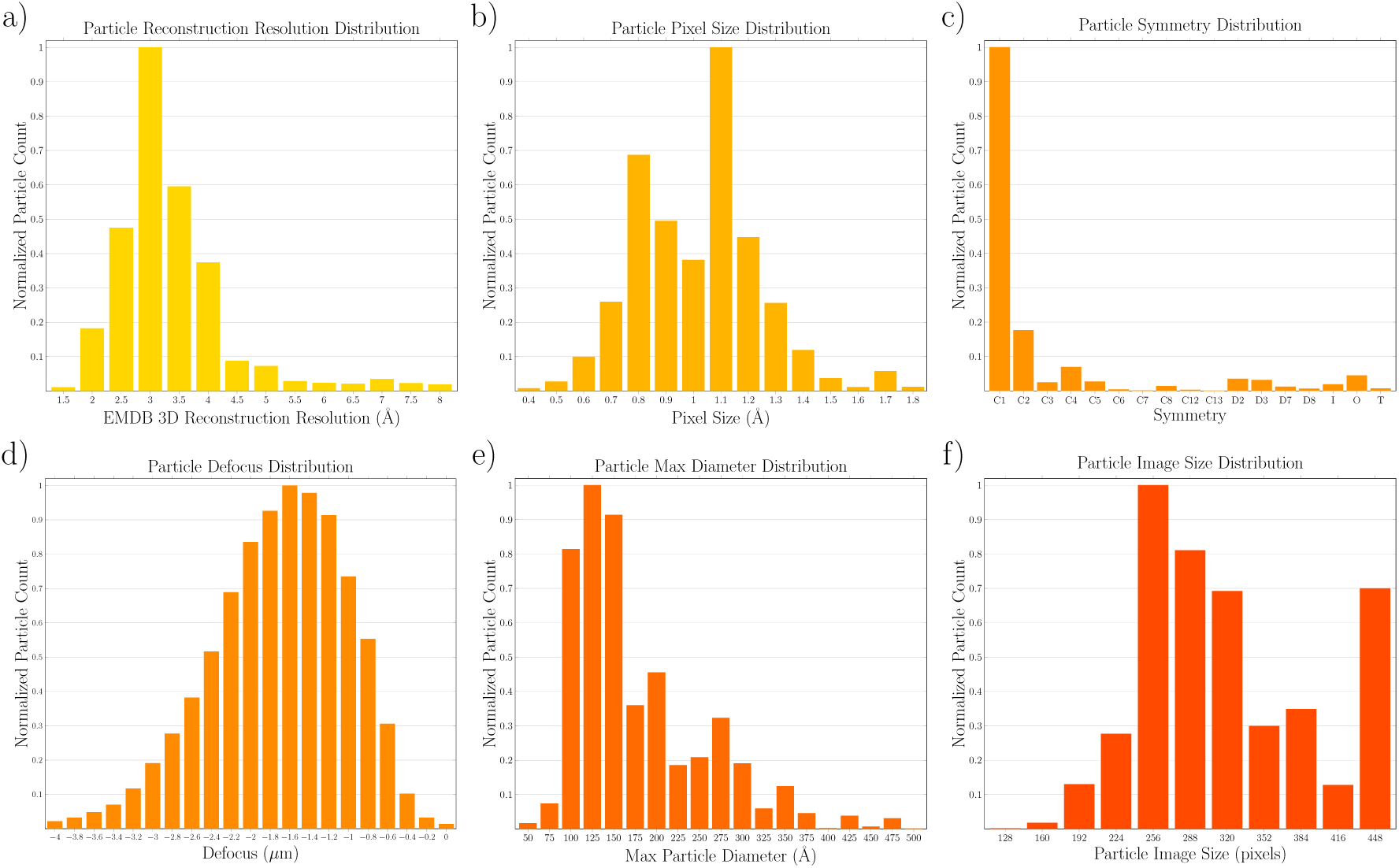
Normalized per-particle distributions of cryoPANDA for: (a) EMDB reconstruction resolution, (b) pixel size, (c) symmetry, (d) defocus values, (e) maximum particle diameter, and (f) particle image size.

### Technical Validation

We validate cryoPANDA along two complementary axes. First, we evaluate the quality of the reconstructions produced by our processing pipeline by comparing them against published reference maps for the same experiments. This assesses whether the curated particle sets retain sufficient information for high-resolution structure determination **(L1)**. Second, we evaluate the dataset’s capacity to support representation learning by training a DINOv2 foundation model from scratch on cryoPANDA particles and probing the learned representations across four downstream tasks: classification of particle properties, micrograph segmentation, particle picking, and particle clustering. These tasks collectively test whether the scale and diversity of cryoPANDA **(L1, L2)**, combined with its rich annotations **(L3)**, enable a single pretrained model to generalize across cryo-EM experiments without task-specific retraining **(L4)**.

### Validating Reconstruction Quality

In Figure 6, we compare the 3D reconstruction resolutions of cryoPANDA and published maps with resolutions better than 6 Å. This is a challenging comparison for cryoPANDA, since in many cases it used only a small fraction of the raw data and consequently reconstructed maps from substantially fewer particles; under otherwise comparable conditions, a larger number of particles generally improves the attainable resolution [286]. Experiments where cryoPANDA improved the resolution are colored green, while those with worse resolution are colored red; in Figure 6 the region within *±*0.5 Å is shaded gray, as these differences are considered modest. In Figure 6a, of the 214 experiments for which cryoPANDA reports reconstruction resolutions, our pipeline yields improved resolution in 75 cases (35%) and inferior resolution in 139 cases (65%). Differences may reflect the use of different subsets of particles for many entries as well as differences in processing software and parameters. For the 139 cases with worse resolution, the mean difference is 0.75 Å. For the 75 cases where cryoPANDA achieves better resolution, the mean improvement is 0.48 Å. In Figure 6b, we explore the correlation between resolution difference (cryoPANDA minus published) against the fraction of particles used by cryoPANDA relative to the published map. As expected, larger resolution differences (exceeding 1 Å) are observed when cryoPANDA used fewer particles (under 100% fraction) than the published reconstruction. In cases where cryoPANDA used more particles than the published map, we observe resolution differences mostly within the *±*0.5 Å.

**Figure 6:**
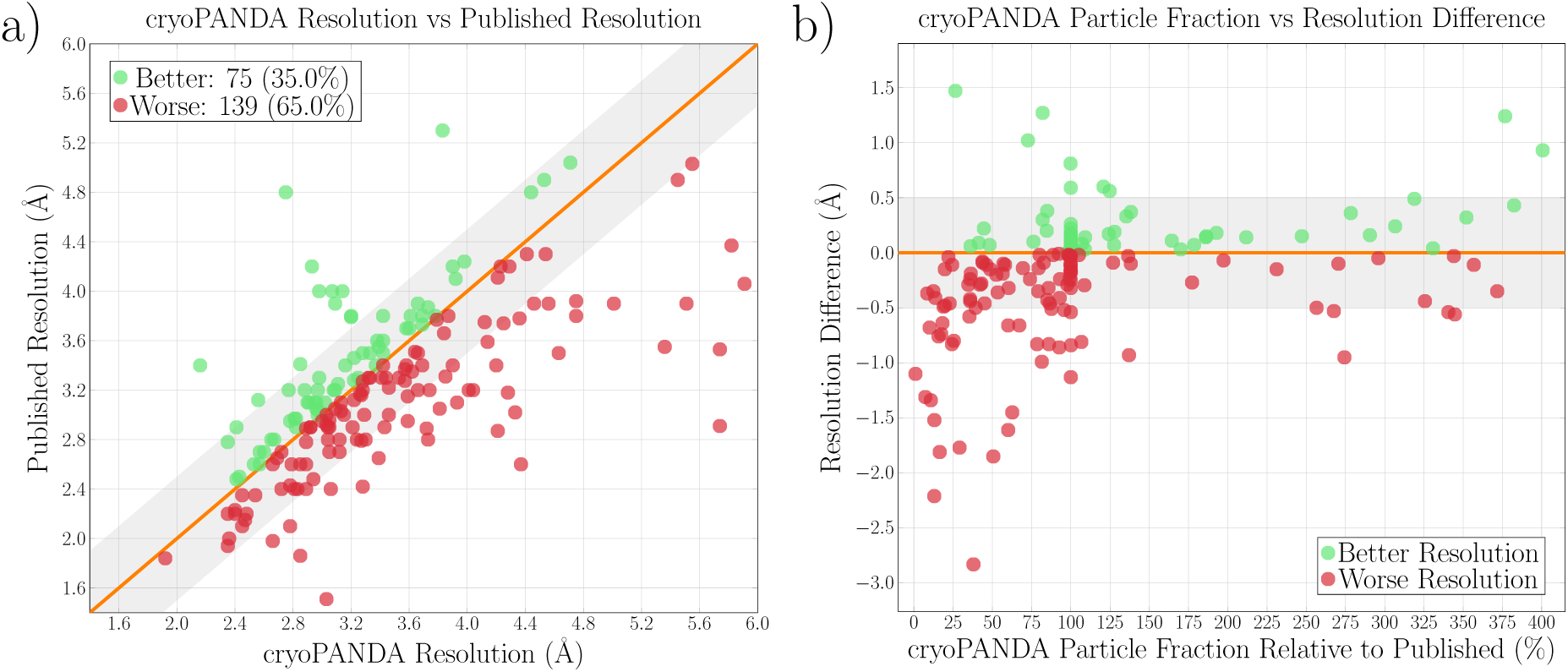
Resolution comparisons between published EMDB maps and those produced by the cryoPANDA pipeline. (a) Reconstruction resolution of cryoPANDA versus EMDB. Points are colored green when cryoPANDA achieves better resolution and red when it yields worse; the region within *±*0.5 Å is shaded gray to denote small differences. (b) Resolution difference (EMDB minus cryoPANDA) versus the particle fraction used by cryoPANDA relative to the published map, using the same color scheme.

An interesting observation is that for many experiments the particle fraction concentrates around 100% on the x-axis; these correspond to EMPIAR entries where we collected all available particles. For these cases, differences in resolution are solely attributed to the *ab-initio* reconstruction and refinement processes steps. Among these entries, cryoPANDA achieves better resolution in 13 cases (mean improvement 0.13 Å) and worse resolution in 8 cases (mean degradation 0.33 Å). Expanding to the 95–105% fraction range, which covers 37 experiments, cryoPANDA reports improved resolution for 17 reconstructions (mean improvement 0.19 Å) and worse resolution for 20 (mean degradation 0.26 Å). These results suggest that the cryoPANDA processing pipeline produces reconstructions broadly comparable to the published maps, especially when using similar particle sets.

In Figure 7, we present five reconstruction examples selected from the 75 experiments for which cryoPANDA achieved better resolution. The figure compares the reconstructed electrostatic potential maps from the published Electron Microscopy Data Bank (EMDB) entries with those from cryoPANDA. The corresponding Fourier Shell Correlation (FSC) plots of the cryoPANDA reconstructions are also shown. The improvements are more noticeable in the first two reconstructions, whereas the remaining three show more modest gains. Across these five examples, the mean improvement is 0.59 Å, compared with 0.48 Å averaged over all 75 improved reconstructions in cryoPANDA.

**Figure 7:**
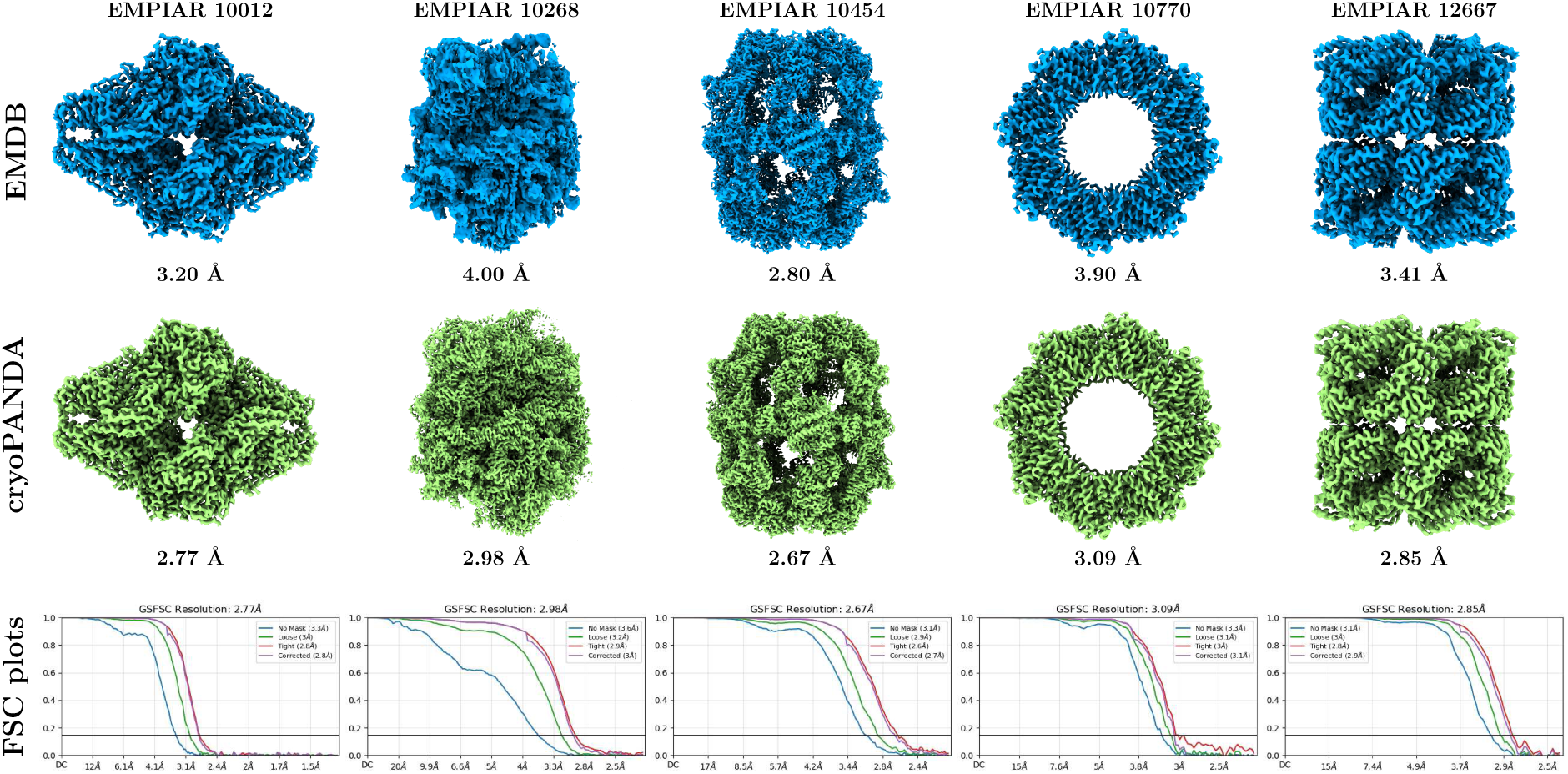
Five reconstruction examples where cryoPANDA achieves better resolution than the published maps. EMDB maps are shown in blue and cryoPANDA density maps in green. The corresponding FSC plots of cryoPANDA are shown below each reconstruction.

### Foundation Model Training

As a proof-of-utility, we demonstrate that cryoPANDA supports representation learning by training a DINOv2 [287] vision foundation model from scratch on the dataset’s particle images. DINOv2 is a self-supervised method that trains a Vision Transformer (ViT) encoder through a student–teacher distillation framework: both networks process augmented crops of each input image, and the student is trained to match the teacher’s output distribution by minimizing their cross-entropy, while the teacher is updated as an exponential moving average of the student (a detailed description of the architecture and training objective is provided in the Supplementary Material). We choose DINOv2 because it represents the state of the art in visual representation learning, and because ViT-based encoders produce both global ([CLS] token) and local (patch-level) representations, enabling evaluation at multiple spatial scales—from full micrographs to individual particle patches.

We adopt the ViT-L/16 configuration from the DINOv2 ImageNet-1K training recipe and partition cryoPANDA into a training set of 215 randomly selected experiments (approx. 32 million particles) and a held-out validation set of 37 experiments (approx. 5 million particles). This experiment-level split ensures that validation evaluations measure generalization to entirely unseen biological samples and imaging conditions, rather than to held-out particles from already-seen experiments. Input particle images, resampled to 224 *×* 224 pixels in the HDF5 distribution, are standardized by subtracting the dataset mean and dividing by the standard deviation. Training runs for 130,000 iterations with a batch size of 2,048 on eight NVIDIA B200 GPUs, completing in approximately 28 hours. The trained teacher encoder is used for all downstream evaluations.

We evaluate the learned representations at two spatial scales. At the micrograph level, we assess whether DINOv2 features can segment particle regions (Evaluation I) and support fully unsupervised particle picking (Evaluation II). At the particle level, we probe whether the representations encode physical and biological properties through linear classification (Evaluation III) and examine how particles from different experiments and complexes organize in the learned feature space (Evaluation IV). For Evaluations I, II, and IV, the encoder is used without any task-specific fine-tuning; for Evaluation III, a single linear head is trained on frozen representations. A schematic of the DINOv2 training scheme applied to cryoPANDA particles is shown in Figure 8.

**Figure 8:**
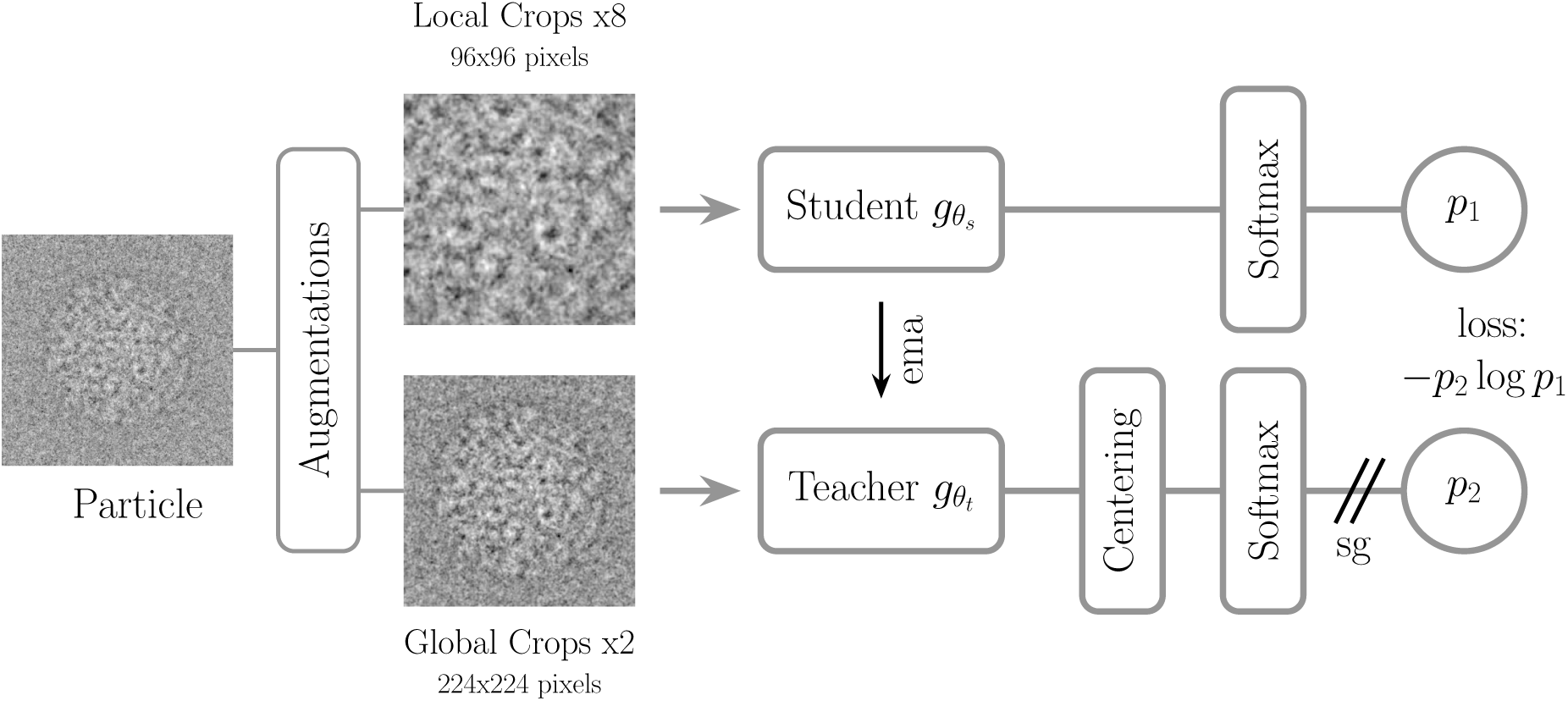
Overview of the self-supervised DINOv2 training scheme. Particle images are augmented into local and global crops, processed by student and teacher encoders, and trained by minimizing the cross-entropy between their output distributions.

#### Evaluation I: Micrograph Level Representations

We assess whether the cryoPANDA-trained DINOv2 encoder produces micrograph-level features that locate particle regions in the micrograph. Since the model is trained exclusively on cropped particle images, this evaluation tests whether the learned representations can be used to segment full micrographs. To extract dense representations, we apply a sliding window of 224*×*224 pixels on each micrograph and extract a feature vector per window from the ViT teacher encoder. Following the visualization protocol of DINO [288], we map the first three principal components of the resulting feature space to RGB channels, where color similarity reflects representational similarity. Figure 9 shows example micrographs alongside their PCA-to-RGB visualizations for nine EMPIAR experiments. Even in low-contrast micrographs such as EMPIAR-10005 [74], EMPIAR-10012 [75], and EMPIAR-12561 [284], particle regions are visually distinct from the background with fine spatial detail, indicating that the encoder separates particle signal from noise without any supervision at the micrograph level. For reference, we also show the micrograph after applying Contrast Limited Adaptive Histogram Equalization (CLAHE), a standard computer-vision technique used to enhance local contrast. In the Supplementary Material, we provide additional segmentation results using single-component PCA and k-means clustering with *k* = 2 (Figures S2 and S3), as well as a PCA-to-RGB segmentation of a tilt-series micrograph acquired during a cryogenic electron tomography experiment (Figure S4), suggesting that a cryoPANDA-trained foundation model can generalize out of the box to cellular imaging modalities such as cryo-ET.

**Figure 9:**
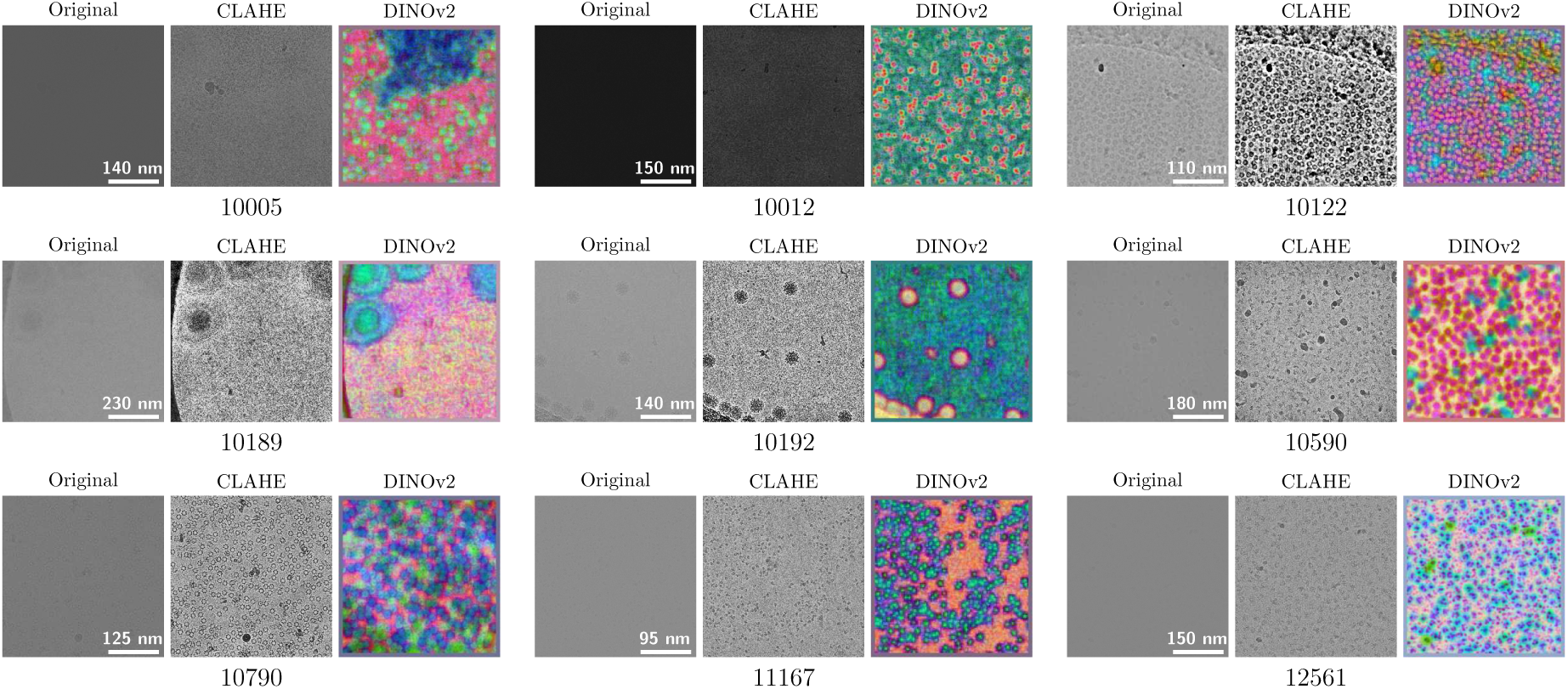
Micrograph-level DINOv2 feature representations visualized by mapping the first three PCA components to RGB. For each EMPIAR experiment, the original micrograph, a CLAHE-enhanced version for improved contrast, and the corresponding DINOv2 PCA representation are shown.

#### Evaluation II: Particle Picking from Micrographs

The micrograph representation results from Evaluation I suggest that the cryoPANDA-trained DINOv2 features carry sufficient spatial information to localize particles. We test this directly by designing a fully unsupervised particle picking pipeline built on top of the frozen DINOv2 encoder, and evaluate it on EMPIAR-10017 [76], which provides manually picked particle annotations for 84 micrographs by Richard Henderson, enabling quantitative comparison. Notably, EMPIAR-10017 belongs to the held-out validation split; therefore, the DINOv2 model has not observed particles from this experiment during training.

### Utilizing Particle Representations for Picking

The picking procedure operates as follows. We partition each micrograph into overlapping 200*×*200 windows (approximately twice the maximum particle diameter) with a stride of 22 pixels, and pass all windows through the cryoPANDA-trained DINOv2 encoder to extract a feature vector per window. For EMPIAR-10017, each micrograph has size 4096 *×* 4096; we extract 31,329 windows per micrograph, yielding a dense representation map of size 177 *×* 177. We then fit PCA on a subset of these representations (156,645 windows from five micrographs) and project all features onto the first principal component, yielding a single scalar score per window. After normalizing these scores to [0, 1] per micrograph, we threshold the resulting score map to identify high-response regions and cluster spatially adjacent detections using topological clustering [289]. Finally, we map detections back to micrograph coordinates and suppress duplicates by merging picks closer than 33 pixels (approximately one sixth of the maximum particle diameter), retaining the higher-scoring detection. One manual step is required: determining which end of the score range corresponds to particle regions, which we resolve by visual inspection of a single example map. A schematic of the pipeline is shown in Figure 10a.

**Figure 10:**
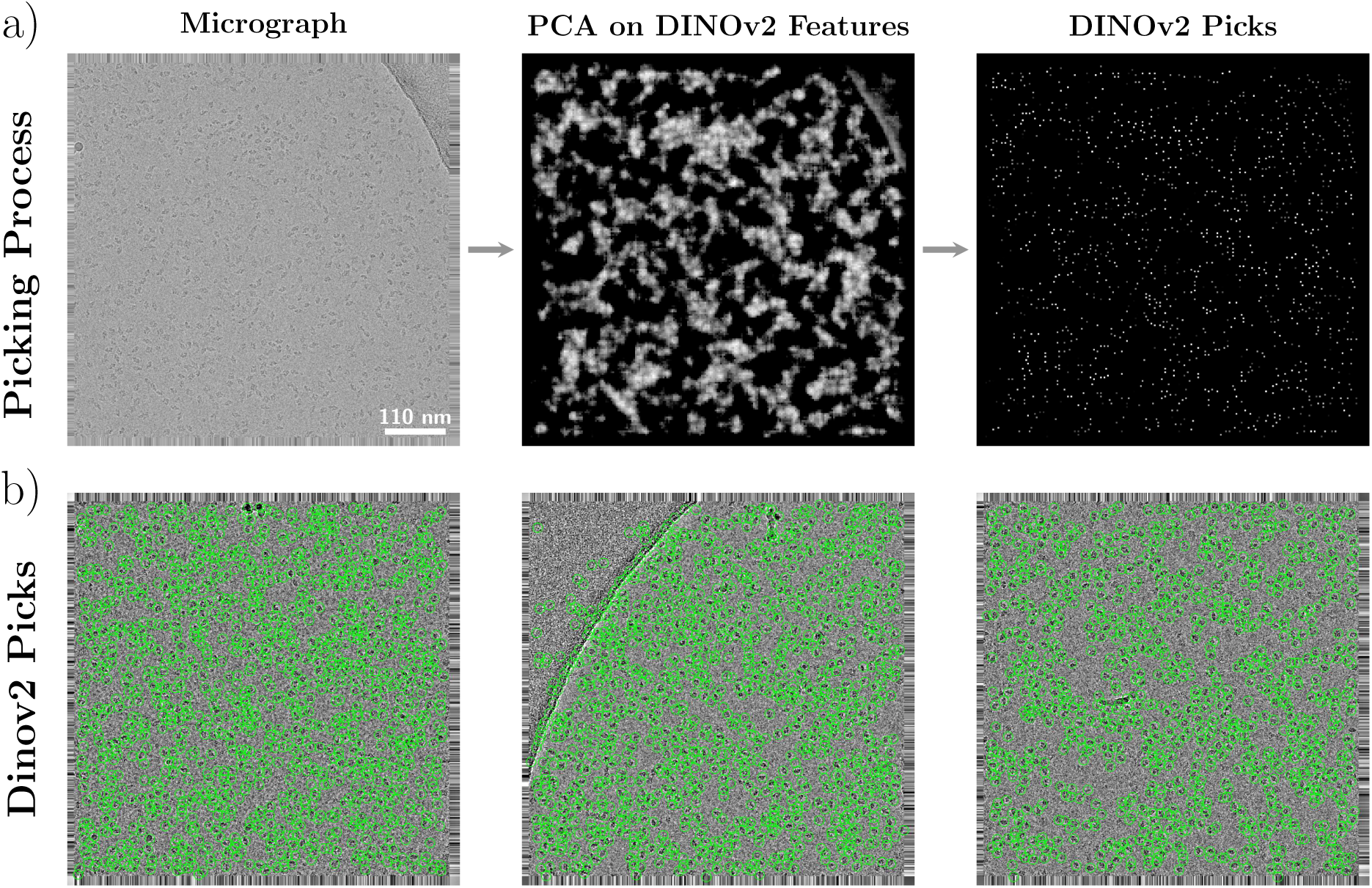
Particle picking based on DINOv2 features. (a) Schematic overview of the DINOv2-based particle-picking workflow. (b) Three example micrographs from EMPIAR-10017, with particle picks identified by the proposed DINOv2 method.

### Particle Picking Performance

We evaluate the DINOv2 picks against Henderson’s annotations. A predicted pick is considered a true positive if the 256 *×* 256 window — the extraction box size used in cryoSPARC for EMPIAR-10017 — centered at the predicted coordinate overlaps by at least 60% with the corresponding window centered at the Henderson annotation. Table 2 reports the results. The DINOv2 picks achieve 91.5% recall and 45.5% precision, yielding an F1 score of 60.8%. The high recall indicates that the *model identifies the vast majority of Henderson’s particles*. The lower precision reflects a more permissive picking strategy — the pipeline selects 78,661 particles compared to Henderson’s 42,755.

**Table 2:** Evaluation of DINOv2-based particle picking on EMPIAR-10017 against Henderson’s manual annotations. Metrics report<u>ed: IoU, Recall, Precision, and F1 score.</u>

| IoU | Recall | Precision | F1 |
| --- | --- | --- | --- |
| 78.9 | 91.5 | 45.5 | 60.8 |

However, a portion of the additional picks may represent valid particles that manual annotation excluded, rather than true false positives. For comparison, the blob picking procedure used during the construction of cryoPANDA yields 185,573 particles from the same dataset, indicating that **the DINOv2-based approach is substantially more selective and can simplify downstream processing** by reducing the burden of pruning and subsequent classification rounds. Picks on three example micrographs are shown in Figure 10b.

### 3D Reconstruction of Picked Particles

To assess whether the DINOv2 picks support downstream structure determination, we perform a 3D reconstruction pipeline using these particles. After two rounds of 2D classification and duplicate removal in cryoSPARC, 43,984 particles remain. Figure 11a compares the resulting reconstruction with the published EMDB map (EMD-2824; 42,755 particles; 4.20 Å) and the cryoPANDA reconstruction (54,270 particles; 4.29 Å). The DINOv2 reconstruction reaches 4.38 Å — within 0.2 Å of both references despite using a fully unsupervised picking procedure. Qualitatively, the three maps are similar; the published map shows marginally finer detail, consistent with its better reported resolution.

**Figure 11:**
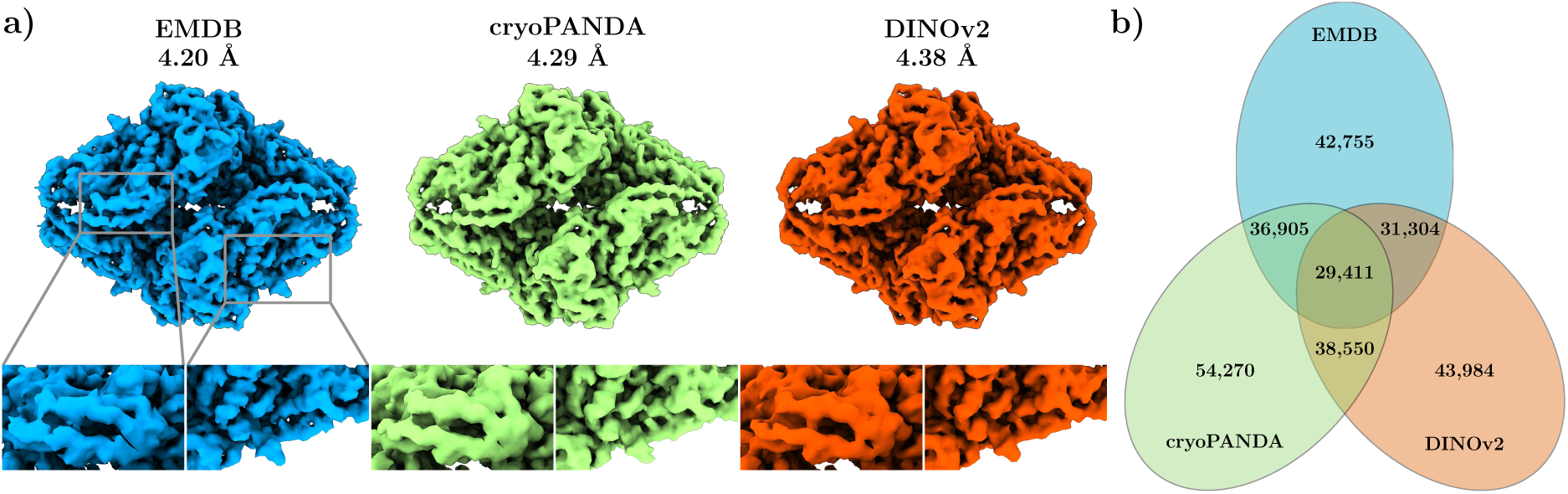
Reconstruction comparisons between published, cryoPANDA, and DINOv2 density maps. (a) Electrostatic potential maps with magnified views of two regions of *β*-galactosidase. (b) Venn diagram showing the overlap between the three particle sets used for the reconstructions.

To understand the relationship between the three particle sets, we quantify their overlap in Figure 11b by matching picks whose centers fall within 50 pixels (half the maximum diameter of *β*-galactosidase, corresponding to 90 Å at 1.77 Å/pixel). We cross-check this distance-based matching against the IoU-based criterion used in Table 2 and obtain very similar overlap estimates. Of the 54,270 cryoPANDA particles, 36,905 (68.0%) overlap with Henderson’s picks, while 31,304 of the 43,984 DINOv2 particles (71.8%) do. The overlap between cryoPANDA and DINOv2 is larger (38,550 particles in common), but the three-way intersection drops to 29,411, indicating that each picking method captures particles the other two miss.

#### Evaluation III: Classifying Particle Characteristics

We assess whether the cryoPANDA-trained DINOv2 representations encode physical and biological properties of particles by training linear classifiers on frozen features. The rich per-particle annotations in cryoPANDA (**L3**) make this evaluation possible: we select six properties spanning acquisition parameters (pixel size, defocus), reconstruction outcome (EMDB resolution), and biological characteristics (symmetry, molecular weight, and maximum diameter). Continuous properties are discretized into equal-width bins. Reconstruction resolution prediction is particularly challenging, as it requires inferring a particle-set-level reconstruction quality metric from a single particle image, making it a difficult task that tests the limits of the representations.

We freeze the pretrained DINOv2 encoder and train a separate linear classifier for each task on frozen features. To disentangle in-distribution performance from cross-experiment generalization, we construct three particle sets: (a) a training set and (b) an in-distribution validation set, both drawn from the 215 DINOv2 pretraining experiments, and (c) an out-of-distribution (OOD) validation set drawn from the 37 held-out experiments unseen during both DINOv2 pretraining and linear probe training. We evaluate under two settings: (i) train on (a), evaluate on (b); and (ii) train on (a), evaluate on (c).

Table 3 reports the results. In setting (i), accuracy exceeds 85% for most tasks (mean 88.4%), demonstrating that the representations are highly linearly separable with respect to these properties. Defocus is the sole exception, reaching approximately 78% — still well above its 25% random baseline, but notably lower than the other tasks. This likely reflects the fact that defocus varies continuously within each experiment and is less correlated with visual particle appearance. In setting (ii), mean accuracy drops to 66.1%, a gap of 22.3 percentage points (pp) relative to setting (i), though still substantially above per-task chance baselines across all six tasks.

**Table 3:** Linear probing accuracy (%) across six classification tasks. **(i)** In-distribution and **(ii)** out-of-distribution (OOD) settings use frozen DINOv2 features; **(iii, iv)** repeat <u>these</u> settings after regressing out acquisition parameters from the features using training-set coefficients only. Random accuracy, 1*/K*, where *K* is the number of classes, is also shown for reference.

| Classification Task | (i) ID | (ii) OOD | (iii) ID Regressed | (iv) OOD Regressed | Random Accuracy |
| --- | --- | --- | --- | --- | --- |
| Symmetry | 93.4 | 60.9 | 94.3 | 61.9 | 5.9 |
| Defocus | 77.6 | 68.2 | 79.8 | 73.1 | 25.0 |
| Max Diameter | 93.8 | 76.9 | 94.3 | 79.9 | 25.0 |
| Molecular Weight | 87.3 | 57.5 | 91.1 | 59.6 | 20.0 |
| Image Pixel Size | 92.4 | 67.7 | 96.4 | 71.5 | 20.0 |
| EMDB Resolution | 86.1 | 65.3 | 88.3 | 67.9 | 25.0 |
| Mean | 88.4 | 66.1 | 90.7 | 69.0 | 20.2 |

To assess whether acquisition-parameter entanglement contributes to the OOD performance gap, we additionally repeat evaluation of settings (i) and (ii) after regressing out the linear influence of experimental acquisition parameters — specifically voltage, electron dose, spherical aberration, amplitude contrast, pixel size and image length — from the frozen features prior to classification. This yields two further settings: (iii) in-distribution and (iv) OOD evaluation on residual features from which the acquisition signal has been removed, with all regression parameters estimated on the training set only and applied without refitting to the validation and OOD sets.

The results, reported in Table 3, reveal a consistent pattern. Removing the acquisition component improves accuracy on *every* task under *both* settings, with mean accuracy rising from 88.4% to 90.7% in-distribution (+2.3 pp) and from 66.1% to 69.0% OOD (+2.9 pp). The largest gains are observed for acquisition-adjacent tasks, particularly defocus prediction, which improves by 4.9 pp in the OOD setting (68.2% → 73.1%). Notably, biologically related tasks also benefit: maximum diameter improves by 3.0 pp under OOD evaluation, and molecular weight by 2.1 pp, indicating that acquisition entanglement was suppressing biologically meaningful signal rather than inflating it.

These results demonstrate that cryoPANDA-trained representations encode a rich set of particle properties with strong linear separability, and that a quantifiable fraction of the cross-experiment generalization gap is attributable to linear acquisition-parameter entanglement rather than a fundamental limit of the learned representations. The per-particle metadata provided by cryoPANDA is sufficient to quantify and partially correct for this entanglement, converting an observed limitation into an interpretable and actionable finding. Closing the remaining gap — for instance through pretraining with randomized acquisition conditions — is a well-motivated direction for future work that cryoPANDA is uniquely positioned to support.

#### Evaluation IV: Particle Level Representations

##### Per-Patch Particle Representations

We examine how the cryoPANDA-trained DINOv2 encoder represents individual particle images. Unlike the micrograph-level analysis in Evaluations I and II, which uses a single feature vector per sliding window, here we extract features per image patch from the ViT teacher encoder, producing representations with a spatial resolution of 16 *×* 16 pixels that reveal structure within each particle. Figure 12 visualizes these per-patch features by mapping the first three PCA components to RGB. Even in low-contrast images, the representations clearly separate the particle from the surrounding background. Beyond this coarse separation, the model captures finer structural detail: for the human adenovirus from EMPIAR-10455 [160], DINOv2 distinguishes the side view of the nucleocapsid from the central core; for EMPIAR-10397 [147], it separates the curved membrane from the aqueous interior; and for FocA [66], the D2 symmetry of the particle is clearly visible in the PCA map. Note that the color coding is determined by PCA initialization and is not directly interpretable.

**Figure 12:**
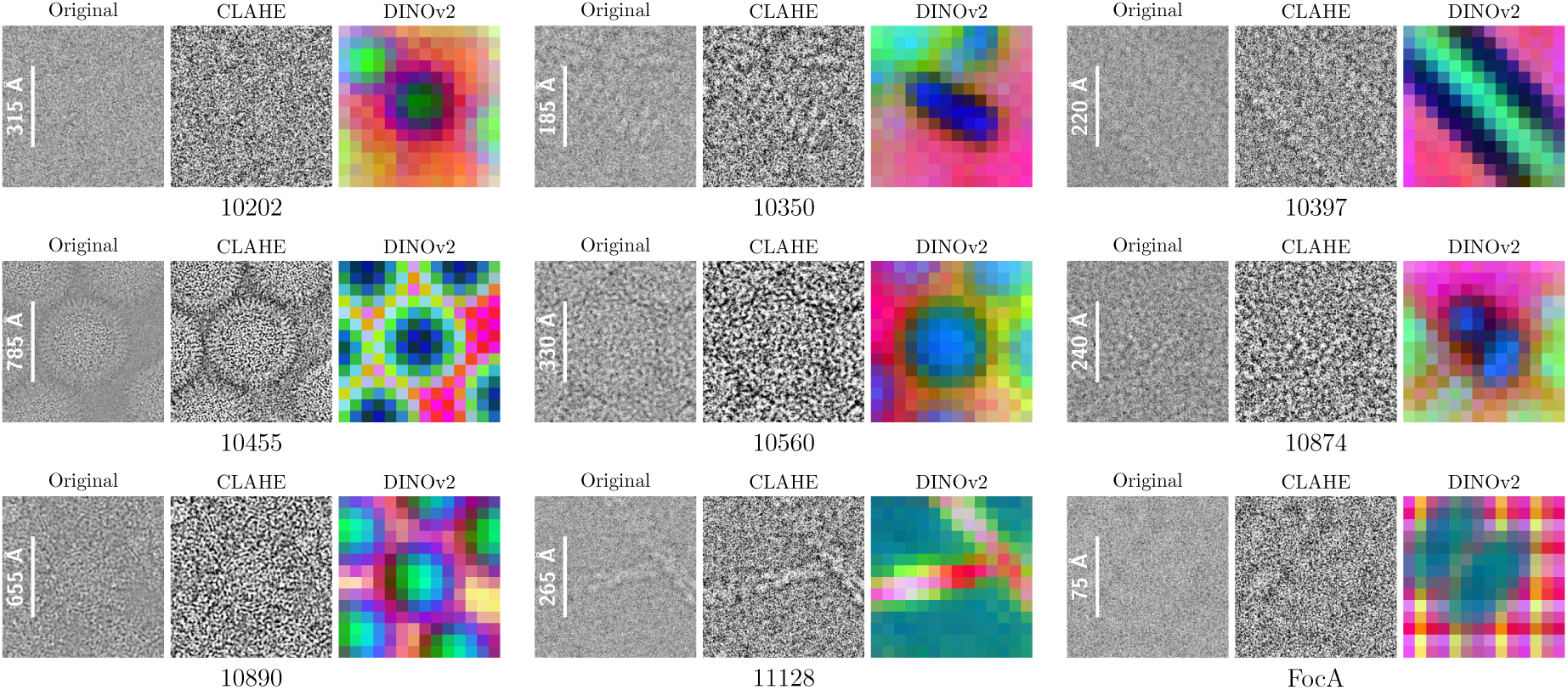
Particle-level DINOv2 per-patch feature representations visualized by mapping the first three PCA components to RGB. For each EMPIAR experiment, the original particle with embedded scale bar, a CLAHE-enhanced version for improved contrast, and the corresponding DINOv2 PCA representation are shown.

##### Cross-Experiment Particle Clustering

We next examine how particles from different experiments organize in the learned feature space. We randomly sample 1,000 particles from each of 30 EMPIAR experiments, separately for the training and validation subsets, and project the DINOv2 features to two dimensions using UMAP [290], as shown in Figure 13.. Across both the validation set (a) and the training set (b), particles cluster strongly by experiment, with only limited overlap between clusters. This experiment-level separation is the dominant structure in the embedding space, whereas microscope parameters and protein characteristics do not appear to align directly with the two main UMAP axes.

**Figure 13:**
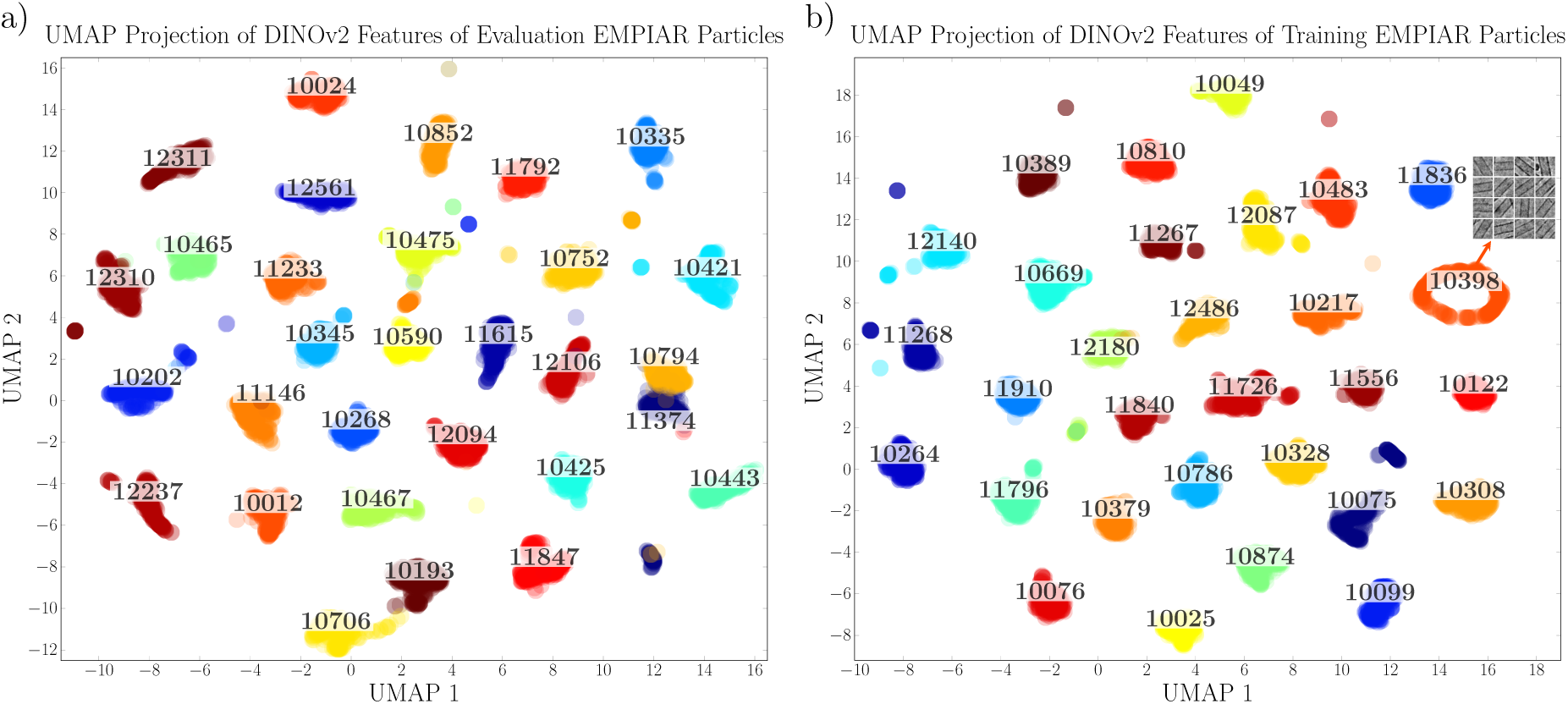
UMAP projections of DINOv2 features for particles from different EMPIAR experiments, shown separately for the (a) evaluation and (b) training sets.

Within this experiment-level clustering, two finer patterns emerge. First, some cluster proximity reflects shared biology: EMPIAR-10193 [108] and EMPIAR-10706 [178], both viral particles, form neighboring clusters, and four ribosome datasets (EMPIAR-11146 [216], EMPIAR-10268 [118], EMPIAR-12094 [270], EMPIAR-10467 [162]) form neighboring clusters in the center of the plot. Second, some proximity reflects shared acquisition conditions rather than biology: EMPIAR-10794 [190] (SARS-CoV-2 NSP15) and EMPIAR-11374 [231] (EXT1-EXT2) are structurally unrelated but cluster together, sharing identical microscope, detector, and optical parameters, as well as nearly identical pixel size, electron dose, and defocus ranges. This observation is consistent with the OOD classification gap reported in Evaluation III — the representations encode acquisition conditions alongside biological features, which can confound cross-experiment generalization. An additional notable observation is the EMPIAR-10398 [147] cluster in subplot (b), which forms a toroidal shape reflecting the diverse in-plane orientations of membrane tubes within this dataset.

##### Intra-Experiment Orientation Separation

We further investigate whether the cryoPANDA-trained DINOv2 representations encode particle orientation information within individual experiments. Figure 14 shows UMAP projections of the DINOv2 features for four EMPIAR datasets, color-coded by dominant viewing direction. Viewing directions are determined by applying Euler angle rotations (ZYZ convention) from the 3D reconstruction to the beam axis and assigning each particle to one of the two most frequent orientations. In all four cases, the two orientation groups form distinct clusters in the UMAP projections, which we validate through separate 2D classification of each cluster in cryoSPARC. In Figure 14d, we observe stronger overlap between the two orientation clusters, which results in shared classes appearing across both 2D classifications — an effect not seen in the other three examples. This separation is not evident in all experiments tested; we present cases where it is most pronounced, and improving its consistency is a direction for future work.

**Figure 14:**
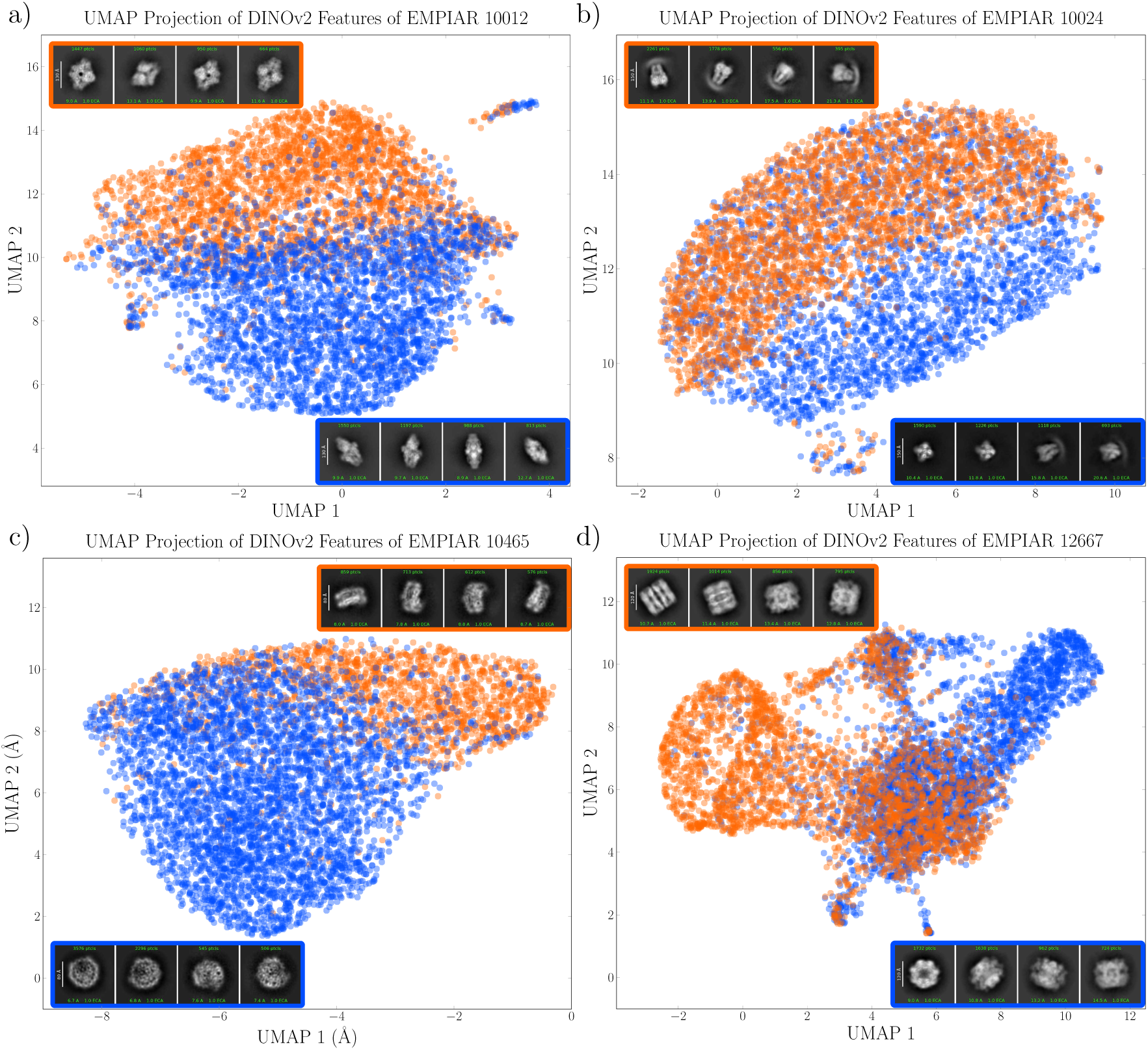
UMAP projections of DINOv2 features for particles from four EMPIAR experiments, revealing two distinct orientation clusters colored by dominant viewing direction. The cluster assignments are validated by subsequent 2D classification of each cluster with cryoSPARC.

##### Intra-Experiment Complex Separation

The strong experiment-level clustering raises a practically important question: can the model separate different macromolecular complexes within a single heterogeneous experiment, where acquisition conditions are shared? To test this, we analyze EMPIAR-10892 [53], a cell extract dataset not included in cryoPANDA, containing four complexes: the 60S pre-ribosome, fatty acid synthase (FAS), the pyruvate dehydrogenase complex E2 core (PDHc), and the oxoglutarate dehydrogenase complex E2 core (OGDHc). We sample 1,000 particles per complex, extract DINOv2 features, and project them with UMAP (Figure 15). PDHc separates strongly from the remaining complexes. The ribosome, OGDHc, and FAS form distinct but neighboring clusters, consistent with their more similar particle dimensions. This result demonstrates that the cryoPANDA-trained model captures biologically meaningful differences even when acquisition conditions are held constant — a capability directly relevant to the field of visual proteomics, where heterogeneous mixtures must be resolved without prior knowledge of their composition.

**Figure 15:**
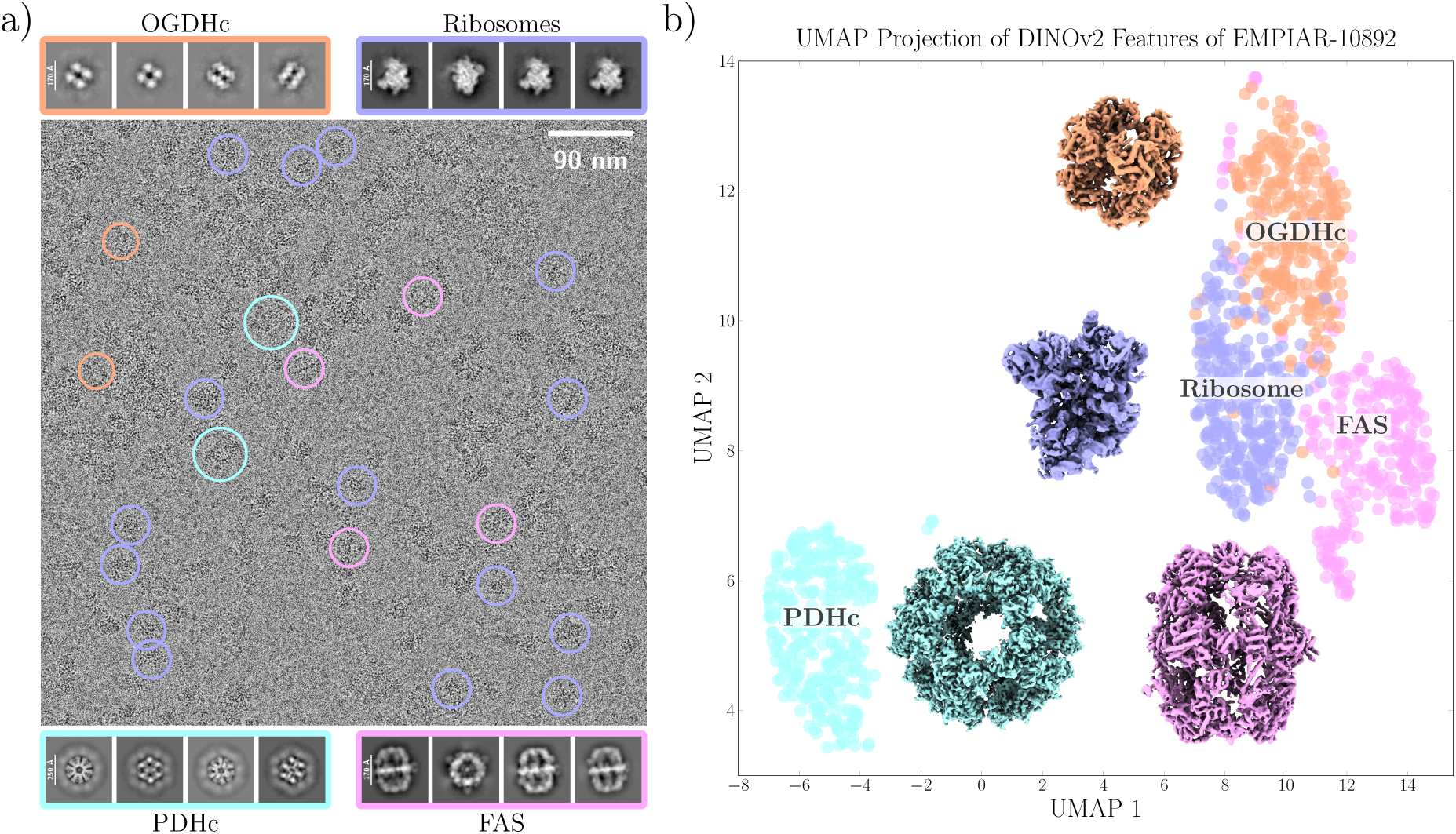
a) An exemplar micrograph from the cell extract experiment EMPIAR-10892, containing particles from four distinct complexes, with representative 2D class averages shown below for each complex. b) UMAP projections of DINOv2 features for particles within the same experiment, with the corresponding 3D reconstruction shown alongside each complex cluster.

The particle-level analysis reveals a clear hierarchy in the learned representations: the model distinguishes between experiments robustly, separates different complexes within heterogeneous mixtures, and in favorable cases resolves dominant viewing orientations within a single complex. Intra-experiment separation remains more challenging than inter-experiment separation, a finding consistent with the OOD classification results of Evaluation III.

Overall, the technical validation empirically demonstrates that cryoPANDA (i) enables the training of large-scale foundation models, which in turn (ii) learn representations that capture semantic information inherent in cryo-EM particles. By training DINOv2 from scratch on over 37 million particles spanning 252 experiments, we show that the resulting model generalizes across diverse downstream tasks — including micrograph segmentation, unsupervised particle picking, linear classification of particle properties, and particle clustering — without any task-specific fine-tuning. cryoPANDA is openly available, together with all processing code and the pre-trained model weights, with the goal of lowering the barrier to developing and benchmarking data-driven methods for cryo-EM analysis. We anticipate that cryoPANDA will facilitate future advances in automated cryo-EM analysis and structural biology.

## Code Availability

All code used for the methodology and technical validation analysis is publicly available at https://github.com/azamanos/cryoPANDA.

## Author Contributions

A.Z., P.K., P.L.K. and Y.P. conceptualized the study. A.Z., F.L.K., and P.K. processed and validated the data. P.L.K. and Y.P. were responsible for the acquisition of funding and resources. All authors contributed to writing and reviewing the manuscript.

