## Supplementary Information for "A 37-million-particle dataset from over 250 experiments to accelerate data-driven cryo-EM analysis"

### Number of Particles per Experiment

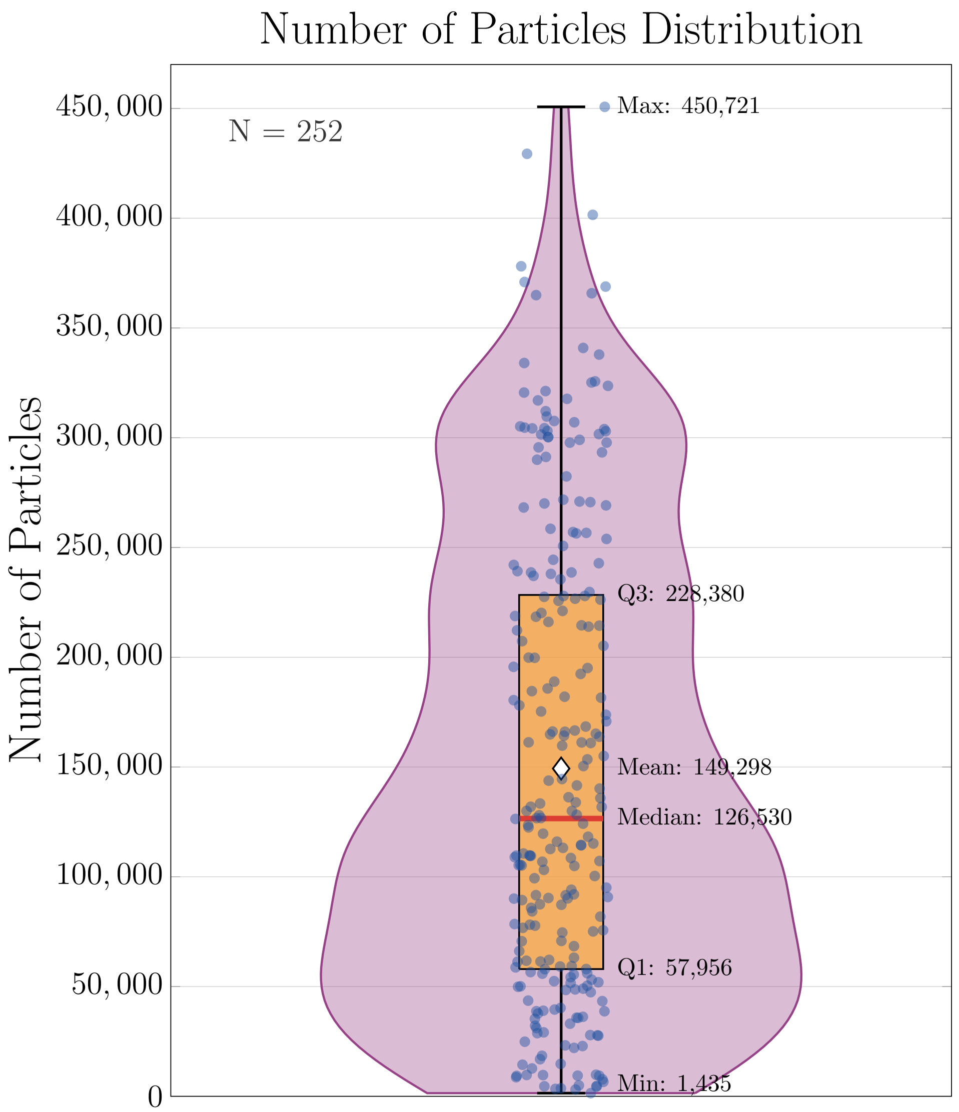


Figure S1: Distribution of particle counts across experiments used in cryoPANDA, shown as a violin plot. Annotated statistics include the mean, median, minimum, maximum, and first and third quartile (Q1, Q3) values.

### DINOv2 Architecture

We briefly summarize the DINOv2 architecture [1]. A schematic overview is provided in Figure 8 in the main text. During training, images in DINOv2 undergo two types of augmentation, the first produces 8 local crops of size $96 \times 96$ pixels, and the second produces 2 global crops of size $224 \times 224$ pixels. These crops are processed through two architecturally identical Vision Transformer (ViT) encoders as follows: all views (both global and local) pass through the student network $g_{\theta_s}$, while only the two global views pass through the teacher network $g_{\theta_t}$. The student's parameters are updated via backpropagation, while the teacher's parameters are updated using an exponential moving average (ema) of the student's parameters:

$$\theta_t \leftarrow \lambda \theta_t + (1 - \lambda)\theta_s, \quad (1)$$

where $\theta_t$ and $\theta_s$ denote the teacher and student parameters, respectively, and $\lambda \in [0.994, 1)$ is the ema decay parameter.

The teacher encoder is the one used for inference at the end of training. During training, the teacher's output representations also undergo centering, a technique that prevents any single dimension of the latent space from dominating. Centering is implemented by subtracting a bias term $c$ from the teacher's output representations before applying the softmax:

$$\tilde{g}_{\theta_t}(x) = g_{\theta_t}(x) - c \tag{2}$$

Centering also encourages teacher's latent representation towards a more uniform distribution. The bias term $c$ is updated as follows:

$$c \leftarrow mc + (1-m)\frac{1}{B}\sum_{i=1}^{B} g_{\theta_t}(x_i), \tag{3}$$

where $m \in [0, 1)$ is a momentum parameter and $B$ is the batch size.

The training objective of DINOv2 is to minimize the cross-entropy between the student and teacher output probability distributions over $K$ prototypes, where $K$ denotes the output dimension of the DINOv2 head, with $K = 65{,}536$ in our setup. These probability distributions are obtained by normalizing the network outputs $g$ with a softmax function. The student distribution is defined as:

$$P_s(x)^{(i)} = \frac{\exp\big(g_{\theta_s}(x)^{(i)}/\tau_s\big)}{\sum_{k=1}^{K}\exp\Big(g_{\theta_s}(x)^{(k)}/\tau_s\Big)}, \tag{4}$$

where $i$ and $k$ denote indices of the output vector $g_{\theta_s}(x)$ and $\tau_s > 0$ controls the sharpness of the distribution. Equivalently,

$$P_s(x) = \mathrm{softmax}\left(\frac{g_{\theta_s}(x)}{\tau_s}\right)$$

Finally, cross-entropy is computed for each pair of global and local views and summed over all such pairs:

$$\min_{\theta_s} \sum_{x\in\{x_1^g, x_2^g\}} \sum_{\substack{x'\in V \\ x'\neq x}} H(P_t(x), P_s(x')) \tag{5}$$

where $H$ denotes the cross-entropy loss between teacher distribution $P_t(x)$ and student distribution $P_s(x')$.

$$H(a, b) = -a \log b$$

The first summation is over the two global views, and the second over all local views in $V$.

#### Micrograph Level Representations

We apply k-means clustering ($k = 2$) to DINOv2 features extracted from a micrograph in EMPIAR-10028 [2] to assess segmentation capability. The results are shown in Figure S2. The two clusters correspond well to particles and background: in the k-means mask shown in (c), the white cluster largely overlaps with particle locations across the micrograph. Subplots (a) and (b) show PCA and UMAP projections of the same features, respectively, and both indicate clear separation between the two clusters. Cluster identities are assigned through visual comparison of the original micrograph and the k-means mask. We highlight EMPIAR-10028 [2] because its particles are clearly visible, making it a useful visual reference.

In Figure S3, we observe consistent particle segmentation across micrographs from multiple EMPIAR experiments [3–14]. These results suggest that the cryoPANDA-trained representations preserve sufficient spatial information to support particle localization, a hypothesis tested directly in Evaluation II of the main manuscript.

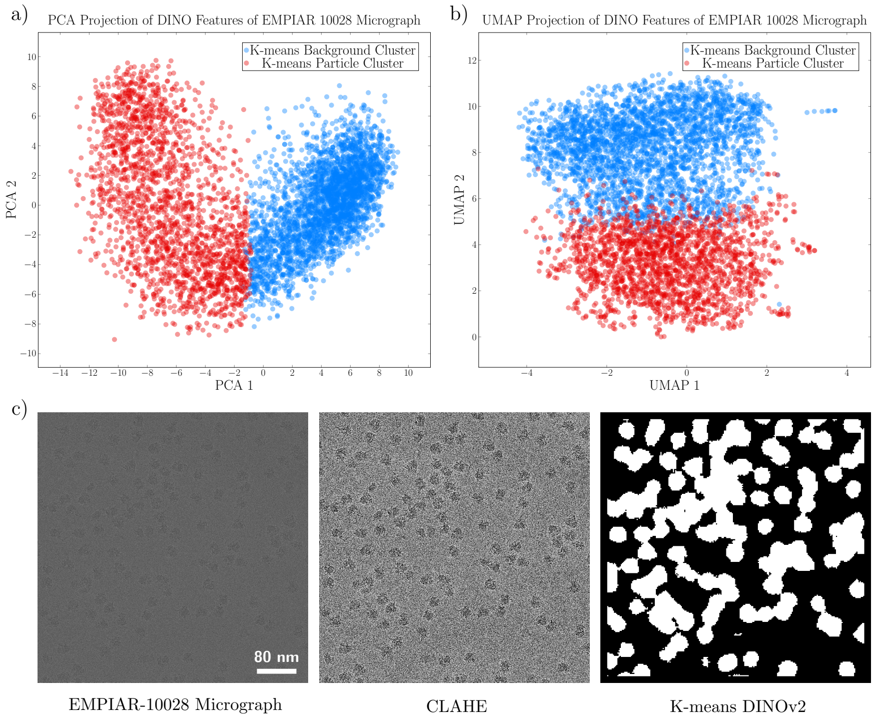


Figure S2: Micrograph-level DINOv2 feature clustering. Subplots (a) and (b) show two-dimensional PCA and UMAP projections, respectively, of DINOv2 features for the micrograph of EMPIAR-10028 [2] in (c), colored by the k-means clustering with two centers. (c) shows the original micrograph, the CLAHE-enhanced version, and the resulting k-means mask, where the two clusters are rendered as white (particle regions) and black (background).

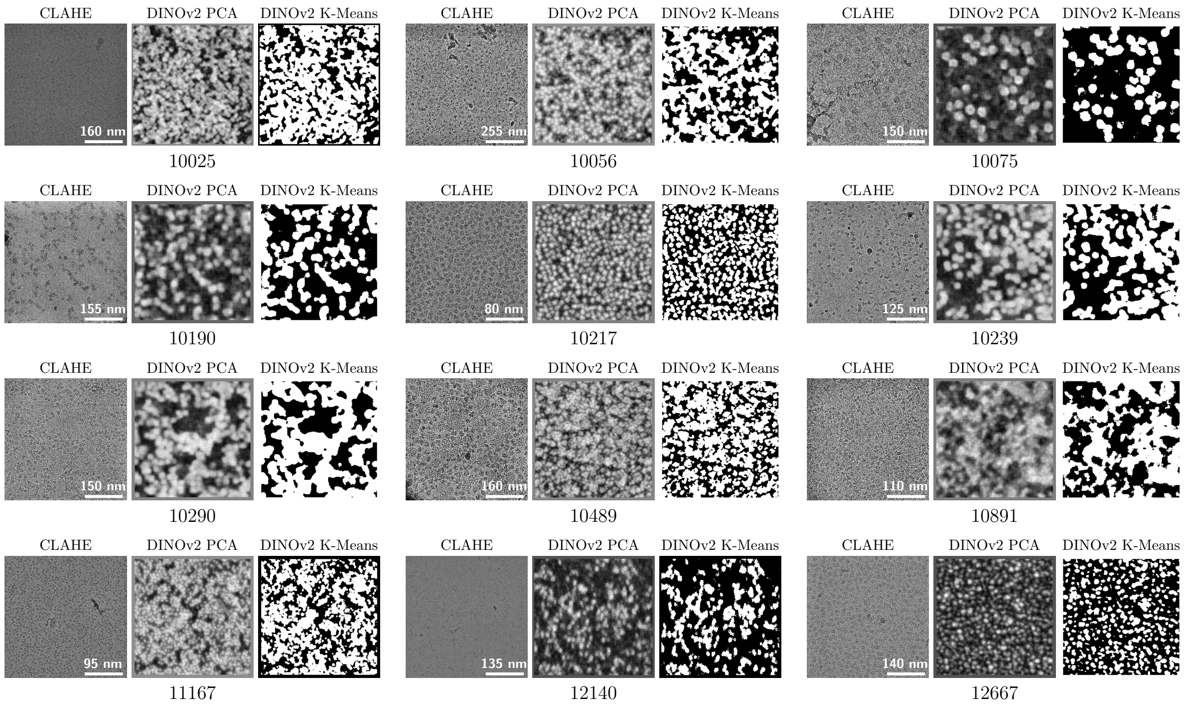


Figure S3: Micrograph-level DINOv2 feature segmentation using single-component PCA (grayscale) and k-means clustering $k = 2$. For each EMPIAR experiment [3–14], the CLAHE-enhanced micrograph, the PCA segmentation, and the k-means segmentation are shown.

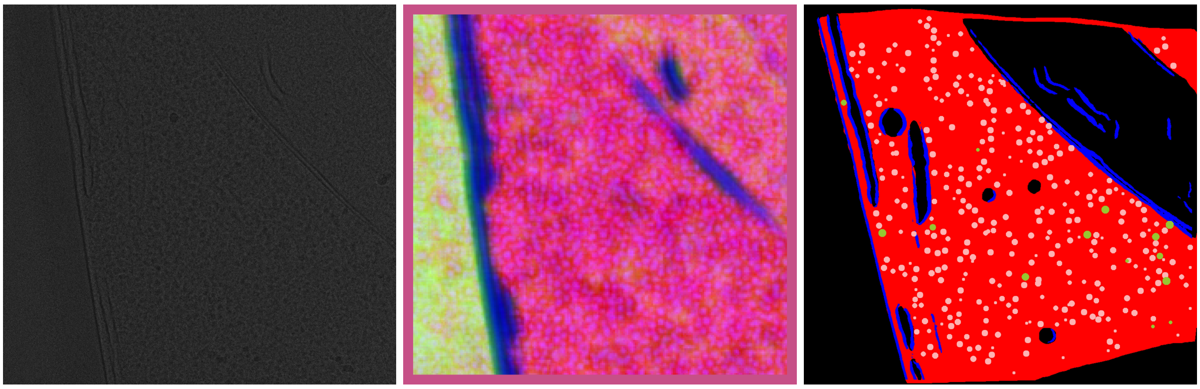

Figure S4: DINOv2 feature representations visualized by mapping the first three PCA components to RGB for a tilt series from EMPIAR-10988 [15]. Left: original tomogram; center: DINOv2 PCA-to-RGB visualization; right: ground truth segmentation mask of the cryo-ET reconstruction. In the segmentation, membranes are shown in blue, cytosol in red, ribosomes in pink, and fatty acid synthase (FAS) in green.
